# Quantitative profiling of lysosomal accumulation through label-free biomarkers via High-Content Holo-Tomographic Flow Cytometry

**DOI:** 10.1101/2025.03.11.642563

**Authors:** Daniele Pirone, Michela Schiavo, Giusy Giugliano, Sandro Montefusco, Lisa Miccio, Pasquale Memmolo, Diego Luis Medina, Pietro Ferraro

## Abstract

Lysosomal storage diseases (LSDs) are genetic disorders caused by enzyme deficiencies that lead to lysosomal dysfunction and progressive cell damage. Accurate visualization and quantification of lysosomes are essential for understanding disease progression and developing effective therapies. Here, for the first time, we successfully identified and characterized lysosomes using an innovative Holo-Tomographic Flow Cytometry (HTFC) technique, which allows label-free, high-content, and high-throughput 3D imaging of lysosomal compartments in single live cells. This breakthrough could revolutionize traditional gold-standard methods overcoming the actual limitations. Leveraging this technology, we discovered novel biomarkers of lysosomal accumulation in LSD-affected cells. In fact, by generating refractive index tomograms, we achieved accurate measurement and comprehensive 3D visualization of cytoplasmic lysosomal aggregation in suspended single cells. Through experimental validation and advanced computational analyses, we identified a quantitative correlation between the 3D lysosomal architecture and the efficacy of various therapeutic strategies, including genetic and pharmacological interventions. This work represents a significant advance in lysosomal research, paving the way for improved diagnostics and the development of targeted therapies for LSDs.

## Introduction

Lysosomal storage diseases (LSDs) encompass over 60 monogenic disorders, mostly inherited in an autosomal recessive manner. They result from defects in lysosomal proteins, including soluble acidic hydrolases, membrane proteins, lipids, enzyme modifiers, activators, or essential non-lysosomal proteins, leading to metabolite accumulation within lysosomes that compromise their integrity and function. This buildup triggers cytotoxic cascades, causing cellular damage, cell death, and ultimately organ dysfunction and degeneration, in many cases affecting the central nervous system (CNS) [1]. Niemann-Pick type C (NPC) disease is a progressive lysosomal lipid storage disorder characterized by a variety of clinical symptoms, including visceral, neurological, and psychiatric manifestations [2,3]. It arises from mutations in the NPC1 gene, a key regulator of intracellular lipid homeostasis [4]. Loss of NPC1 function leads to the intracellular accumulation of various lipid species, such as cholesterol, glycosphingolipids, sphingomyelin, and sphingosine primarily impacting the CNS, liver, and spleen [2,3,5,6]. While there is no cure for NPC, treatments such as *Miglustat* and *Cyclodextrin* aim to manage symptoms and slow disease progression [7,8]. Unfortunately, a lack of tools to assess the drug’s in-situ operation at the single-cell level hampers our understanding of its mechanism of action and its limitations, hindering efforts to optimize its efficacy and minimize adverse effects.

NPC diagnosis combines clinical evaluation, biochemical and genetic tests, and imaging to detect cholesterol accumulation and lysosomal dysfunction. However, lysosomal physical properties such as morphology, surface characteristics, density, and homogeneity have not been deeply studied in health and disease yet, despite they might influence physiology in health and disease conditions. Various imaging techniques, including 3D fluorescence microscopy, enable the visualization of intracellular organelles such as lysosomes [9]. However, despite their significance in cell biology, these methods have certain limitations, including the need for extensive sample preparation and the requirement for high-quality and selective fluorescently conjugated probes or selective antibodies [10]. These analyses involve sample fixation and staining, which can introduce artifacts that alter the native cellular structures. Moreover, most of the above techniques have limitations as they analyze adherent cells. In fact, physical constraints from flat substrates can distort organelle organization, causing lysosomal aggregates and even the nucleus to appear clustered or misshapen. Similarly, Transmission Electron Microscopy (TEM) provides high-resolution images but requires thin sectioning, extensive preparation, trained personnel, and costly equipment [11]. More broadly, many advanced imaging techniques such as high content imagers [12] are limited by high instrumentation costs [13].

Recently, a 2D study based on label-free Quantitative Phase Imaging (QPI) [10,14-17] was reported detecting lysosomal localization in the perinuclear area in wild type (WT) and NPC1 knockout (KO) cells [18], demonstrating its potential as a diagnostic tool for NPC and drug evaluation. Building on this preliminary result, here we show for the first time that Holo-Tomographic Flow Cytometry (HTFC) [19] can provide high-content imaging of lysosomal aggregates in single live suspended cells. Indeed, we demonstrate that it is possible to retrieve detailed volumetric and accurate biophysical information by a label-free, high-throughput, 3D imaging, thus avoiding the physical constraints in adherent cells [20-22]. In this study, we leverage HTFC for discovering and demonstrating new biomarkers [23] extracted from 3D refractive index (RI) tomograms in statistically significant large populations of cells (∼2000). We show that this approach provides unique advantages for characterizing lysosomal morphology, which could improve disease diagnosis and efficient screening of genes or small molecules that modulate lysosomal aggregation.

Here, we collect and analyze 3D data from WT and NPC1 KO HeLa cells, leveraging pharmacological and genetic treatments to modulate the lysosomal compartment. This approach provides insights into lysosomal identification, disease mechanisms, and potential therapeutic strategies in NPC1-deficient cells. To induce NPC-like lysosomal storage, we use the cholesterol transport inhibitor U18666A, while transient expression of WT NPC1 in NPC1 KO cells restores lysosomal function (Fig. 1a). Additionally, we evaluate two lysosome-targeting therapeutic strategies, i.e. cyclodextrin-based drug therapy to reduce lipid storage and siRNA-mediated SPAG9 depletion to reduce lysosomal aggregation (Fig. 1b). By using HTFC morphometric analysis to phenotype the lysosomal changes in the different conditions, we establish a unique set of 3D morphometric parameters to full characterize lysosomal alterations in LSD-affected cells, thus marking a significant advancement in lysosomal research at the aim to enhance diagnostics and pave the way for targeted LSD therapies.

**Fig. 1.**
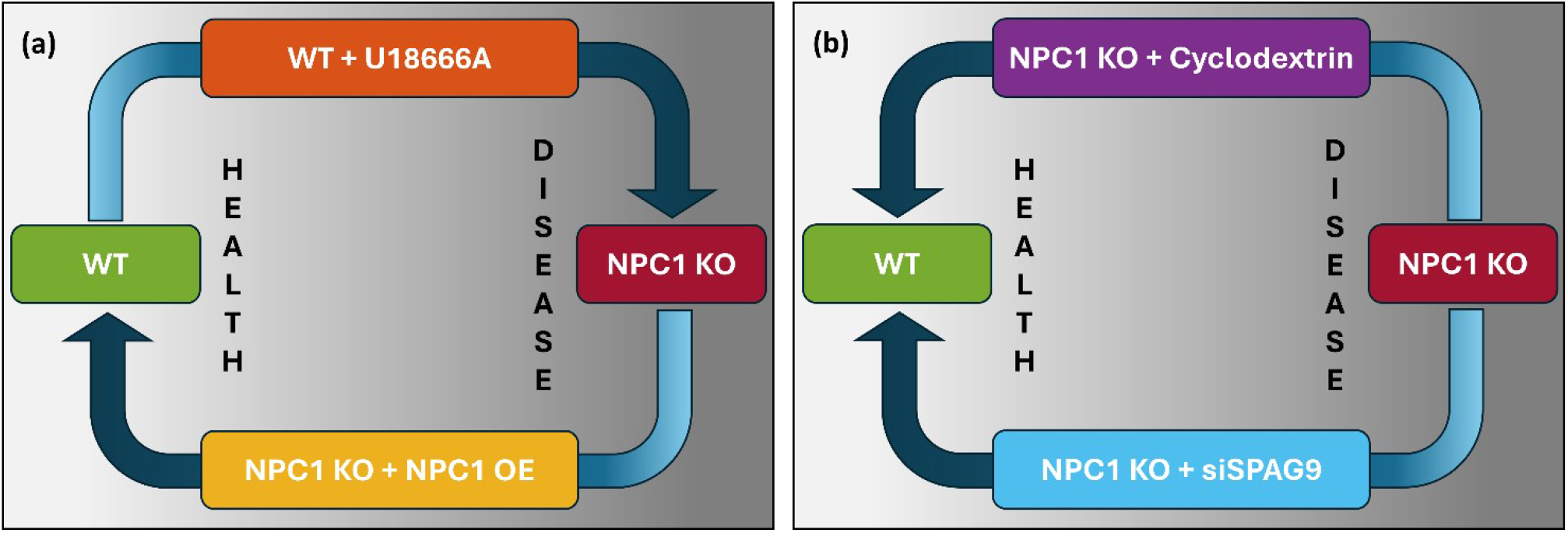
Conceptual scheme to characterize the lysosomal compartment in health and disease conditions by using 3D morphometric parameters established using HTFC. **(a)** The U18666A cholesterol transport inhibitor aims at inducing the NPC phenotype, while the NPC1 gene overexpression (OE) aims at recovering the WT phenotype. **(b)** The cyclodextrin-based drug and the SPAG9 gene silencing (siSPAG9) are examined as possible therapeutic strategies to rescue morphological alteration of lysosomal compartment in HeLa cells from the NPC disease condition.

## Experimental Results and Data Analysis

We experimentally recorded and numerically reconstructed the tomograms with the 3D RI distributions of 1939 HeLa cells flowing in suspension along the microfluidic channel of our HTFC system, as described in the Methods sections. As summarized in Table S1, the entire dataset of HeLa cells was collected in 5 different key experiments, indicated as experiments A, B, C, D, and E. For each experiment, two different cell lines were acquired, as described in the following sections.

### 3D label-free visualization of lysosomal aggregates

We developed a dedicated processing pipeline for the 3D RI tomograms. In fact, to establish a novel method for segmenting the lysosomal compartment in 3D label-free manner using HTFC, we employed two distinct lysosomal morphological conditions in HeLa cells: WT cells and an LSD model lacking the NPC1 gene (i.e., NPC1 KO cells) [2,3,5]. High-content confocal imaging showed an accumulation of the lysosomal compartment towards the perinuclear area in NPC1 KO cells, in contrast to the more uniform distribution observed in WT cells (Fig. S1 and Fig. S2). These pronounced pathological changes in NPC1 KO cells provided an ideal contrast to facilitate the clear identification of the lysosomal compartment using HTFC. Thus, we analyzed all the 358 HeLa WT cells and 766 HeLa NPC1 KO cells collected during the 5 experiments (Table S1). The central slices of the 3D RI tomograms of a WT cell and an NPC1 KO cell are displayed in Figs. 2(a,f), respectively. Holographic Tomography (HT) [24-26] is a label-free imaging technique that uses the RI distribution within cells to identify intracellular structures without staining [14,27]. However, differentiating organelles can be challenging due to overlapping RI values [28,29]. Recent advancements aim to improve specificity in QPI [29,30] and HT [29,31,32]. To address these challenges in the flow cytometry environment of HTFC, where imaging is further complicated by the lack of a substrate and the cell’s suspended condition, a method called Computational Segmentation based on Statistical Inference (CSSI) was developed [20], successfully identifying specific organelles (e.g., nucleus and vacuoles) by analyzing statistical relations among RI patterns [20,21]. Instead, in the easiest case of lipid droplets (LDs), they were segmented through a simple thresholding by exploiting their distinguishable RI values [22].

**Fig. 2.**
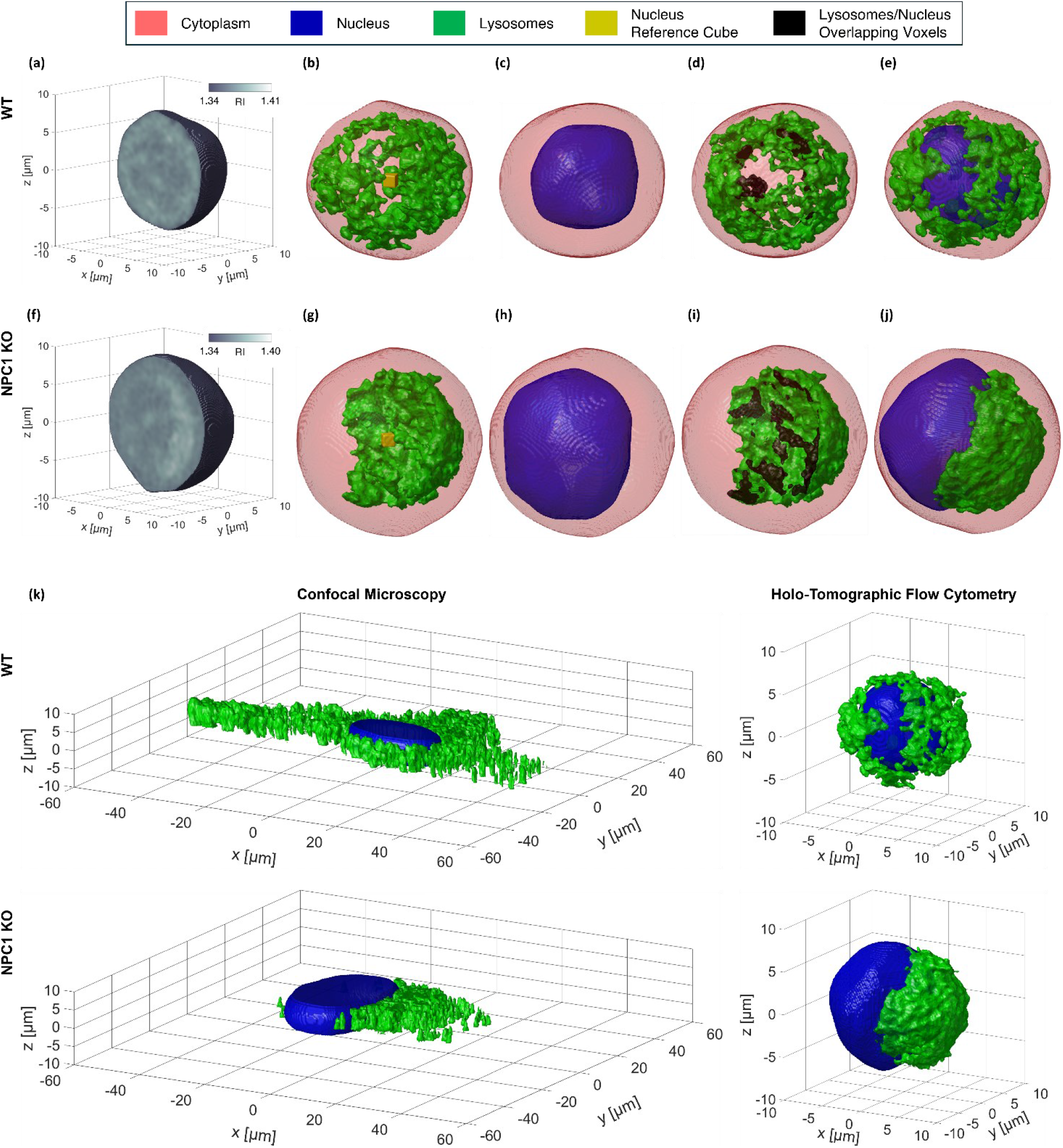
Segmentation of the lysosomal volumes’ container (LVC) in a suspended (a-e) HeLa WT cell and (f-j) HeLa NPC1 KO cell recorded and reconstructed by HTFC. **(a,f)** Central slice of the 3D RI tomogram reconstructed by HTFC. **(b,g)** Threshold-based segmentation of the rough lysosomal volume (green) within the cell shell (red), along with the reference cube (yellow) for the CSSI-based segmentation of the nucleus. **(c,h)** CSSI-based segmentation of the nucleus (blue) within the cell shell (red). **(d,i)** LVC (green) obtained after refining the rough lysosomal volume in (b,g) by removing the volumes (black) overlapped with the nucleus in (c,h), respectively. **(e,j)** Cell shell (red) with the segmented nucleus (blue) and LVC (green). **(k)** On the left, 3D images of lysosomes stained by LAMP1 (green) and nucleus stained by DAPI (blue), obtained by high-content confocal imaging of a HeLa WT cell (top) and a HeLa NPC1 KO cell (bottom). On the right, label-free 3D images of lysosomes (green) and nucleus (blue), obtained by HTFC imaging of a HeLa WT cell (top) and a HeLa NPC1 KO cell (bottom), corresponding to the cells in (e,j), respectively.

Lysosomes are known to have high RI values inside the cell [18,33,34]. Therefore, here we developed a new strategy by combining a suitable RI thresholding and the CSSI method to identify the 3D intracellular lysosomal compartment inside each single-cell RI tomogram, as described in the Methods section and shown in Figs. 2(b-d,g-i). The nuclear and lysosomal compartments segmented in a HeLa WT cell and a HeLa NPC1 KO cell can be observed together in Figs. 2(e,j), respectively. An enhanced 3D rendering of Fig. 2 is displayed in Supplementary Movie S1 and Supplementary Movie S2, where the entire slice-by-slice visualization of RI tomograms in Fig. 2(a) and Fig. 2(f) and the full rotation of binary tomograms in Figs. 2(b-e) and Figs. 2(g-j) are reported for the HeLa WT cell and the HeLa NPC1 KO cell, respectively. In agreement with the health and disease phenotype observed by fluorescent-based high-content confocal analysis (Fig. S1 and Fig. S2), label-free HTFC revealed uniformly distributed lysosomes in the cytoplasm of WT cells, whereas NPC1 KO cells showed a dramatic aggregation of the lysosomal compartment in the perinuclear area covering a pole of the adjacent nucleus. Remarkably, lysosomes, which often form aggregates, are small organelles that can be difficult to identify individually, even with advanced microscopy techniques like HTFC. Therefore, we will refer to the segmented volume containing one or more lysosomes as a “lysosomal volume container” (LVC).

### Quantitative 3D morphometric biomarkers in NPC healthy and diseased cells

In addition to the label-free advantage, a visual comparison between the 3D images from a high-content confocal microscope and the label-free HTFC segmentations in Fig. 2(k) indicates that 3D HTFC data from suspended cells provide more reliable morphometric information than 3D confocal data from adherent cells. This can be seen much better through direct comparison between the two imaging tools in Supplementary Movie S3 and Supplementary Movie S4 containing the same HeLa WT and HeLa NPC1 KO cells, respectively. In fact, when cells are in suspension, the 3D arrangement of lysosomes relative to the nucleus can be more accurately captured, whereas adhesion conditions may introduce distortions by cell spreading on a substrate, potentially affecting the reliability of 3D confocal analysis. HTFC overcomes this limitation by analyzing cells in suspension, ensuring that measurements reflect intrinsic intracellular phenotypes rather than artifacts introduced by adhesion. By providing robust statistical analysis of large cell populations under identical conditions, HTFC offers a complementary approach to traditional gold-standard imaging techniques of adherent cells. The ability to characterize lysosomal morphology in a label-free, high-throughput manner presents a unique opportunity to systematically quantify lysosomal alterations in NPC disease in 3D.

To illustrate this, we identify a novel set of quantitative biomarkers to comprehensively analyze the complete dataset of 358 HeLa WT cells and 766 HeLa NPC1 KO cells gathered using our HTFC system. The 17 quantitative 3D morphometric features, described in the Supplementary Information (Section S1 and Table S2), are sorted in Fig. 3(a) based on their Fisher’s discriminant ratio (FDR) defined in Eq. (S3) [35], which rank them according to their ability in discerning the healthy (WT) or diseased (NPC1 KO) single-cell state. Specifically, we were able to identify three main groups of features (see the different colors in Fig. 3 and the scheme in Fig. S8), termed here as *lysosomes-nucleus biomarker* (it measures the perinuclear arrangement of lysosomal aggregates), *nucleus-cell biomarker* (it is related to how lysosomal accumulation affects the position and shape of the nucleus), and *lysosomes-cell biomarker* (it describes the spatial arrangement of lysosomal aggregates in relation to the cell’s biovolume). As shown in the pie chart of Fig. 3(b), the lysosomes-nucleus arrangement is the most discriminant biomarker (average FDR of 0.55), followed by the lysosomes-cell arrangement (average FDR of 0.41), and the nucleus-cell arrangement (average FDR of 0.18). This reflects the biological knowledge about asymmetric accumulation of lysosomes around the nucleus as a distinctive characteristic of the NPC disease and confirms the HTFC reliability. Another possible categorization of the 3D morphometric features is related to the coordinate reference system in which they are computed, i.e. cartesian and spherical, as underlined by the different symbols in Fig. 3(a) and by the scheme in Fig. S8. As displayed in the pie chart of Fig. 3(c), features in spherical coordinates (average FDR of 0.44) are more distinctive than features in cartesian coordinates (average FDR of 0.34). While this categorization may be irrelevant in adherent cells, it makes much more sense for the quantitative analysis of lysosomal arrangement in suspended cells.

**Fig. 3.**
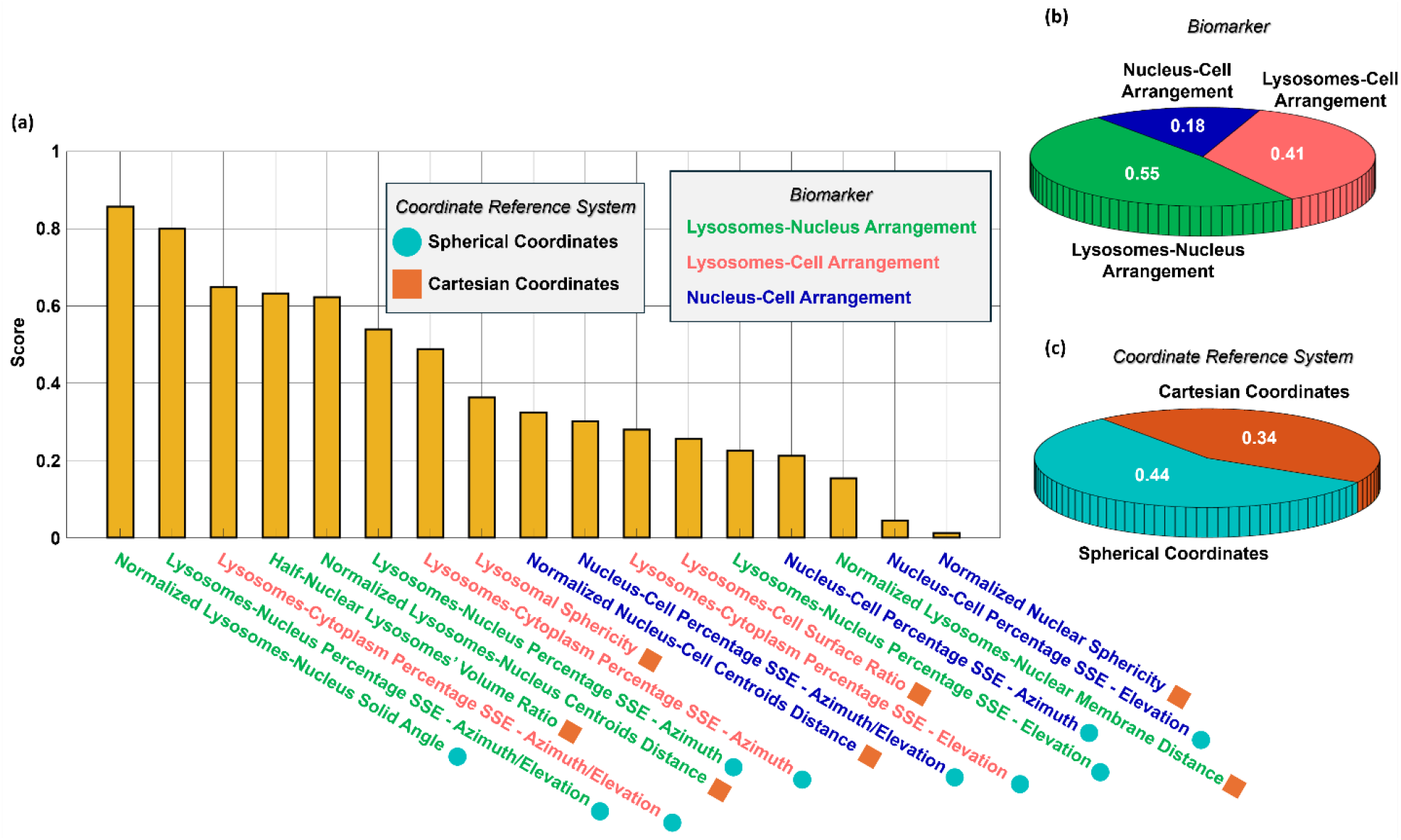
Ranking of the 3D morphometric features computed from the 3D RI tomograms of label-free suspended HeLa WT cells and HeLa NPC1 KO cells to characterize the healthy and diseased state. **(a)** FDR score, used to sort the 17 morphometric features. Features can be computed in a cartesian or a spherical coordinate reference system, and they can be categorized into three classes of biomarkers according to the specific intracellular arrangement they describe. **(b)** Pie chart related to the categorization based on the biomarker. **(c)** Pie chart related to the categorization based on the coordinate reference system. In (b,c), for each class, the average FDR scores of their features are reported.

According to the feature ranking in Fig. 3(a), the normalized lysosomes-nucleus solid angle (NLNSA) appears to be the most distinctive feature of the NPC disease, followed by the lysosomes-nucleus percentage sum of squared error (PSSE) of both the azimuth and elevation coordinates (LNPSSE_Az-El_), lysosomes-cytoplasm PSSE of both the azimuth and elevation coordinates (LCPSSE_Az-El_), and half-nuclear lysosomes volume ratio (HLVR). To better understand the concept behind the quantification of lysosomal asymmetries inside a suspended cell, these features are illustrated in Fig. 4. Moreover, the HLVR and NLNSA features can be also seen in Supplementary Movie S5 and Supplementary Movie S6, respectively, by observing different cells from several perspectives.

**Fig. 4.**
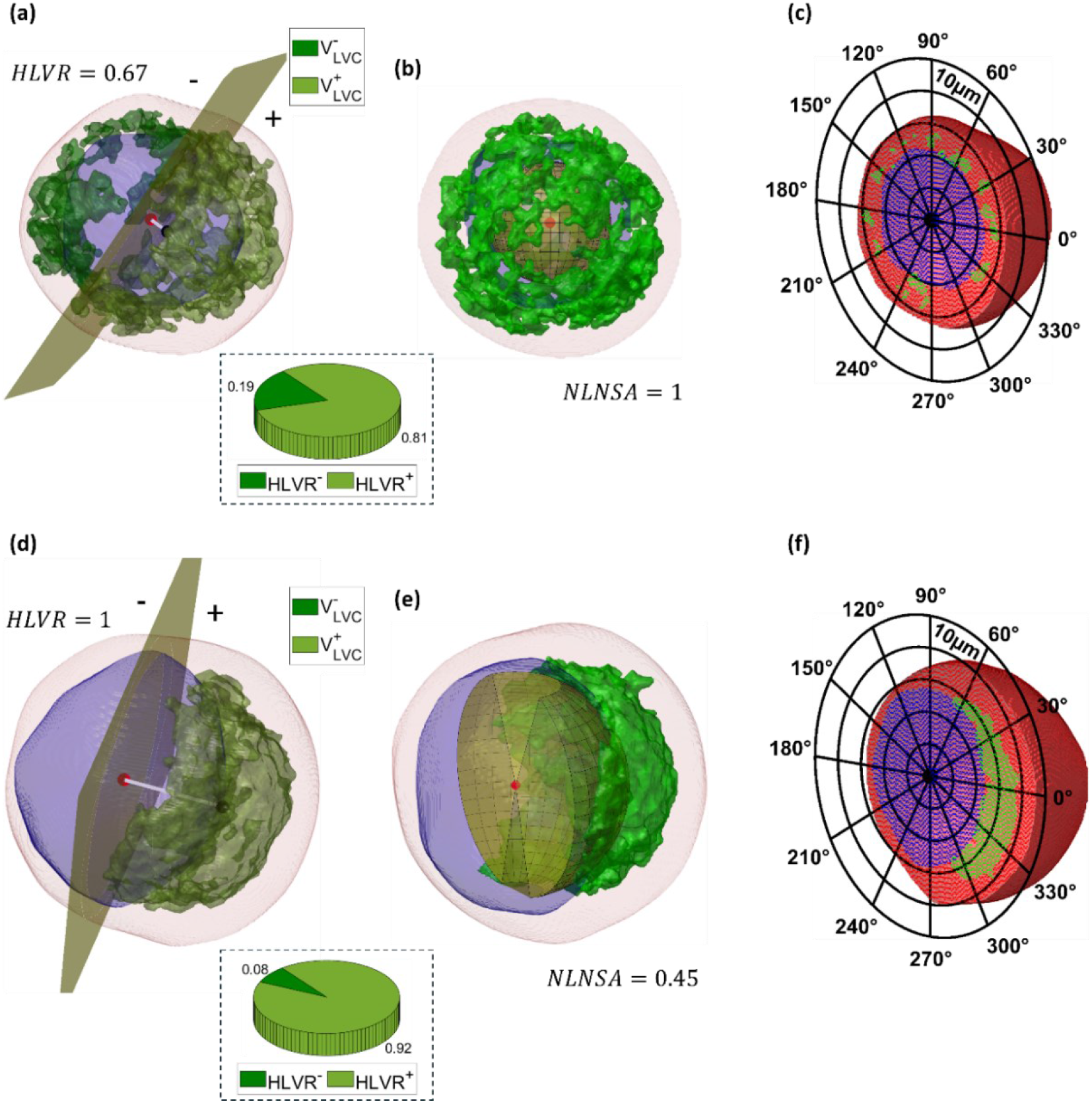
Quantitative morphometric characterization of the lysosomal spatial distribution (green) with respect to the nucleus (blue) in the 3D RI tomogram (red) of a suspended (a-c) HeLa WT cell and (d-f) HeLa NPC1 KO cell. **(a,d)** The cutting plane (yellow) passing through the nucleus center (red dot) and orthogonal to the segment (white line) joining the nucleus centroid (red dot) to the LVC centroid (black dot) divides the LVC compartment into two parts, i.e. one in the half-space (+) containing the LVC centroid (olive green volume), and the other one in the half-space (−) not containing the LVC centroid (dark green volume). The HLVR parameter is defined as the ratio between the LVC volume in the positive half-space and the total LVC volume. It measures how uniformly distributed the lysosomes are around the nucleus. Low values (≥0.5) are related to a uniform distribution of lysosomes around the nucleus, while high values (≤1) are related to a lysosomal accumulation around the nucleus. The HeLa NPC1 KO cell in (d) has a highly asymmetric lysosomal accumulation, which is well quantified by the HLVR with respect to the much more uniform perinuclear distribution of lysosomes in the HeLa WT cell in (a). To further verify this, the pie charts in the insets report the average *HLVR*^−^ and *HLVR*^+^ values over the whole population of 358 HeLa WT cells and 766 HeLa NPC1 KO cells, respectively. The sizes of the two slices of pie are much more unbalanced in the diseased HeLa NPC1 KO cells (d) than the healthy condition of HeLa WT cells (a), thus quantifying the accumulation on one side of the nucleus. The striking asymmetry in lysosomal compartment distribution observed in HeLa NPC1 KO cells (d), compared to the balanced distribution in healthy HeLa WT cells (a), can be considered a quantitative parameter able to assess the progression of NPC disease and potentially serve as an early diagnostic marker. **(b,e)** The NLNSA parameter reported at the bottom measures the solid angle (yellow) that subtends the LVC compartment (green), centered in the centroid (red dot) of the nucleus (blue). It quantifies the lysosomal accumulation with respect to the nucleus. High values (≤1) are related to a uniform distribution of lysosomes around the nucleus, while low values (≥0) are related to a lysosomal accumulation around the nucleus. **(c,f)** Central slice of the 3D RI tomogram after the intracellular segmentation, reported in a polar coordinate system centered in the nucleus centroid (black dot). This allows to foresee how a curvilinear coordinate system (i.e., polar in 2D and spherical in 3D) allows to measure better the relative positions of the several intracellular components inside a quasi-spherical cell, i.e. lysosomes (green), nucleus (blue), and cytoplasm (red). Indeed, by converting the cartesian coordinates of the voxels belonging to the LVC, the nucleus, and the remaining cytoplasm into a spherical coordinate system, an additional set of morphometric features can be introduced to describe the spatial distribution of the LVC compartment in the 3D space of a suspended cell, which are related to the azimuth and elevation coordinates.

### Assessment of the quantitative analysis in 3D label-free suspended cells

To assess the ability of the proposed HTFC feature set in quantitatively evaluating changes in the volumetric behavior of the lysosomal compartment upon both pharmacological and genetic manipulation, we follow the scheme sketched in Fig. 5(a). In experiment B (Table S1), we induced cholesterol accumulation by treating HeLa WT cells with the cholesterol transport-inhibiting compound U18666A, which is widely used to mimic NPC disease [36,37], thus obtaining WT + U18666A cells (Fig. S3), as sketched in the top branch of Fig. 5(a). Conversely, in experiment C (Table S1), to revert disease phenotype such as the perinuclear localization of the lysosomal compartment, we transiently transfected HeLa NPC1 KO cells with a gene overexpression (OE), i.e. with a plasmid encoding a WT version of the NPC1 gene, thus obtaining NPC1 KO + NPC1 OE cells (Fig. S4), as sketched in the bottom branch of Fig. 5(a). High-content confocal analysis confirmed the effects of U18666A increasing the accumulation of cholesterol within the lysosomes in the perinuclear area of WT + U18666A cells (Fig. S3). Instead, the overexpression of WT NPC1 gene in NPC1 KO cells resulted in a partial recovery of the lysosomal positioning from the perinuclear area to a more uniform localization in the cytoplasm (Fig. S4). Indeed, NPC1 OE reduced cholesterol accumulation in the lysosomal compartment, thus reflecting a correction of the NPC1 phenotype (Fig. S5). Tomograms shown in the scheme of Fig. 5(a) provide visual proof of the correct intracellular specificity retrieved by the high-content HTFC label-free imaging and the proposed segmentation method of 3D suspended cells, which agrees with high-content confocal fluorescence imaging.

**Fig. 5.**
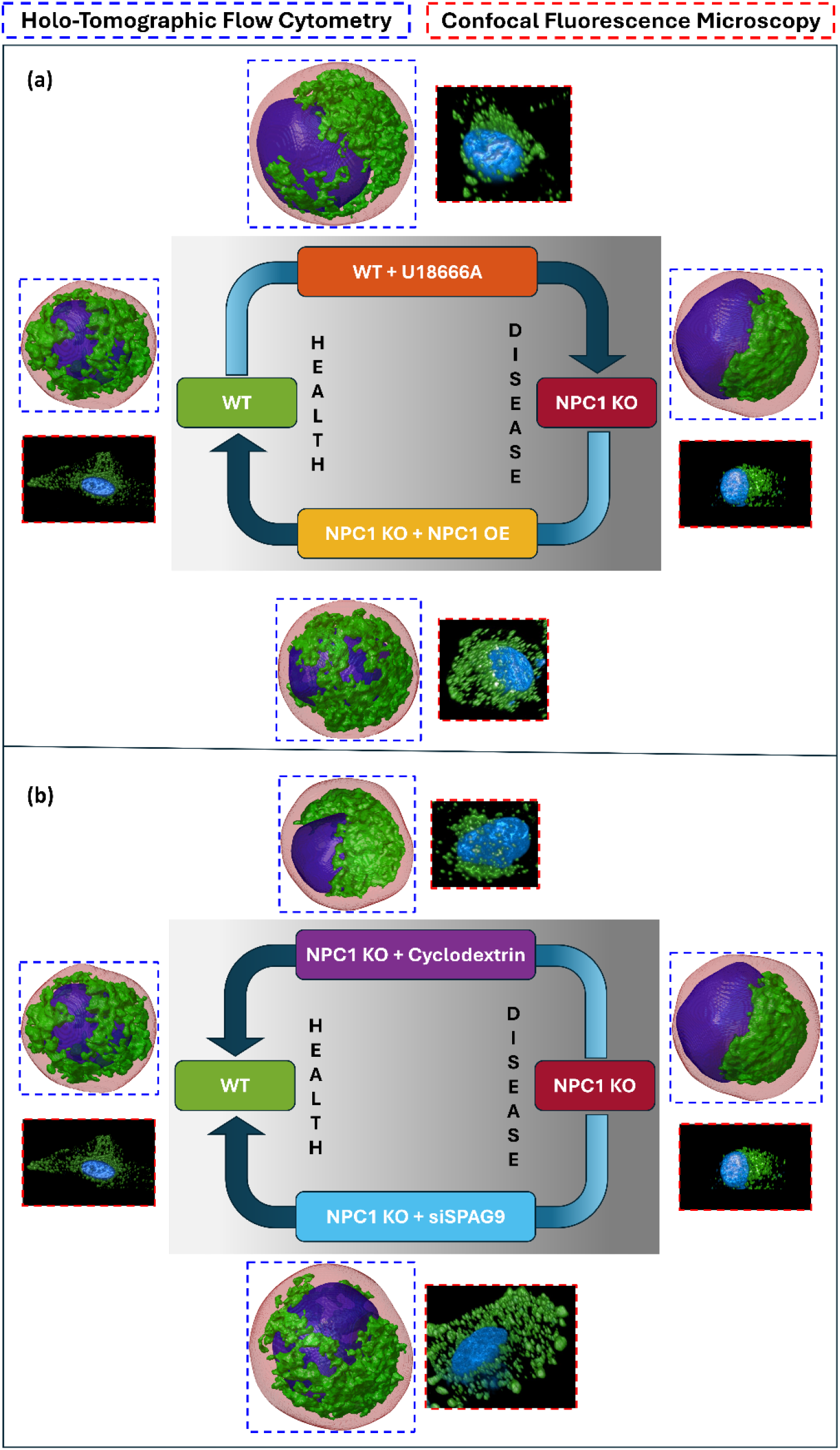
Examples of cells in the NPC health and disease cycle imaged by high-content HTFC (dashed blue) and high-content confocal imaging (dashed red). In label-free HTFC images, the nucleus (blue) and the lysosomal aggregates (green) are segmented inside the whole cell (red). In confocal images, the lysosomal aggregates (green) are marked by LAMP1 staining and the nucleus (red) is marked by DAPI staining. Label-free HTFC imaging of cells in suspension provides a much more reliable snapshot of the actual intracellular lysosomal arrangement in the 3D space than the adherent cells imaged by confocal fluorescence microscopy. **(a)** Assessment cycle. Starting from the WT condition of a uniform lysosomal distribution around the nucleus, the U18666A treatment leads the WT cell to the NPC phenotype with a lysosomal accumulation on one side of the nucleus. Conversely, starting from the NPC condition, the gene overexpression allows the cell to be rescued by the lysosomal accumulation, thus going back to a uniform spatial distribution of the lysosomal compartment. **(b)** Drug testing cycle. Starting from the NPC condition, the cyclodextrin treatment and the SPAG9 depletion aim to rescue the cell by the lysosomal accumulation and positioning, thus going back to a uniform spatial distribution of the lysosomal compartment.

We measured the features herein proposed for each 3D RI tomogram to compare the different lysosomal phenotypes among the several cell lines. We considered as ground truth the entire dataset of 358 HeLa WT cells and 766 HeLa NPC1 KO cells collected during experiments A-E (Table S1). Then, we compared changes observed separately during experiments B-E to the ground truth and each other. In fact, during experiments B-E, HeLa cells have undergone specific biological treatment that might have introduced changes in the lysosomal phenotype of the corresponding control case. We quantified these changes through the percentage variation (PV) defined in Eq. (S4). Let’s consider as example the swarm charts of the NLNSA and HLVR features in Figs. 6(a,b), respectively. NLNSA and HLVR belong to the lysosomes-nucleus biomarker, and they are the highest ranked features computed in spherical and cartesian coordinates, respectively (Fig. 3(a)). Both measure the degree of lysosomal accumulation around the nucleus (lower NLNSA values and higher HLCR values indicate a greater accumulation). As expected by the visual analysis in Fig. 5(a), the two opposite treatments (i.e., U18666A in experiment B and gene OE in experiment C) cause changes with opposite sign in both these two parameters.

**Fig. 6.**
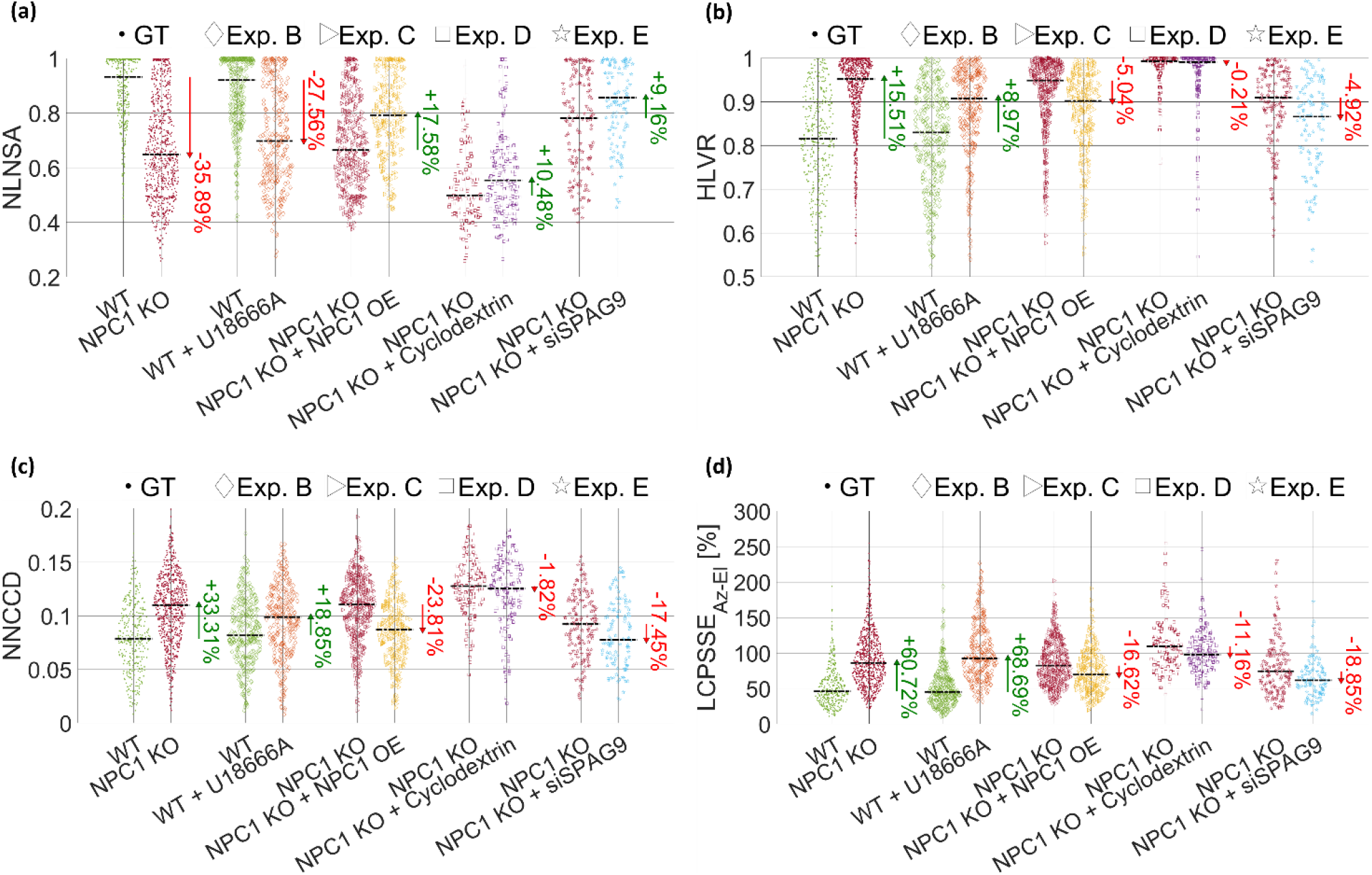
Comparison between the highest ranked features’ swarm charts about the ground truth (GT) experiment (WT vs. NPC1 KO) and other four experiments (B, C, D, and E) based on specific treatments applied to the HeLa cells to quantitatively evaluate the lysosomal compartment by HTFC. For each experiment, the PV values are reported (green if positive, red if negative), computed from the median values (black dashed lines). **(a)** Normalized Lysosomes-Nucleus Solid Angle (NLNSA). By comparing the WT cells to the NPC1 KO cells in the ground truth experiment, the lysosomal accumulation typical of the NPC disease substantially affects the NLNSA feature, which decreases by 35.89% in the NPC1 KO case. In the experiment B, the U18666A treatment applied to WT cells to emulate the NPC disease also leads to a 27.56% decrease in the NLNSA parameter with respect to the control WT case. Instead, the gene overexpression applied in experiment C to the NPC1 KO cells to recover the WT condition leads to a 17.58% increase in the NLNSA parameter with respect to the control NPC1 KO case. In both experiments D and E, the NLNSA parameter has a similar PV passing from the NPC1 KO + Cyclodextrin and NPC1 KO + siSPAG9 to the control WT cells (+10.48% and +9.16%, respectively), opposite in sign to the PV passing from WT cells to NPC1 KO cells (−35.89%). **(b)** Half-nuclear Lysosomes Volume Ratio (HLVR). A positive PV is observed in the ground truth experiment passing from WT cells to NPC1 KO cells (+15.51%) and in the experiment B emulating the NPC condition (+8.97%), while a negative PV is observed in experiment C restoring the WT condition (−5.04%). Moreover, while maintaining the opposite sign with respect to the ground truth, the PV is much lower in the cyclodextrin treatment of experiment D (−0.21%) than the SPAG9 gene silencing of experiment E for the HLVR (−4.92%). In particular, the HLVR PV is almost null in the cyclodextrin case. **(c)** Normalized Nucleus-Cell Centroids Distance (NNCCD). It is greater in the NPC1 KO case of ground truth (+33.31%) and in the WT + U18666A case of experiment B (+18.85%) with respect to the WT population, while it decreases in the NPC1 KO + NPC1 OE case of the rescue experiment C (−23.81%). Moreover, the NPC1 KO + siSPAG9 cells in experiment E have a negative PV of -17.45%, which is much higher in absolute terms than the -1.82% PV of the NPC1 KO + Cyclodextrin in experiment D. **(d)** Lysosomes-Cytoplasm PSSE of both the Azimuth and Elevation coordinates (LCPSSE_Az-El_). It is higher in the NPC1 KO case of the ground truth (+60.72%) and in the WT + U18666A case of experiment B (+68.69%) with respect to the WT population, while it decreases in the NPC1 KO + NPC1 OE case of the rescue experiment C (−16.62%). Moreover, the PV is opposite in sign to the ground truth in both experiments D and E, but, in absolute terms, siSPAG9 has a greater effect than cyclodextrin (−18.85% and -11.16%, respectively).

Another effect of the lysosomal accumulation observed in the 3D RI tomograms is a change of the nuclear position and shape. In the tomograms of Fig. 5(a), nucleus appears more spherical and concentric with the cell in the healthy case (WT) while it becomes more flattened towards the cellular membrane due to the lysosomal accumulation typical of the NPC condition (NPC1 KO). This is well quantified by the normalized nucleus-cell centroids distance (NNCCD) in Fig. 6(c), i.e. the highest ranked feature within the nucleus-cell biomarker, computed in cartesian coordinates (Fig. 3(a)).

Within the lysosomes-cell biomarker, the highest ranked feature is the LCPSSE_Az-El_, which is computed in spherical coordinates (Fig. 3(a)). As reported in Fig. 6(d), in agreement with the expected outcome of ground truth experiment and experiments B-C, higher values of this parameter indicate a greater lysosomal accumulation around the nucleus and a greater decentralization and deformation of the nucleus with respect to the cell centroid and shape.

Hence, label-free HTFC can segment lysosomes and follow their morphological and positioning changes upon pharmacological and genetic manipulations. A discussion about the other proposed morphometric biomarkers is reported in the Supplementary Information (Section S2 and Figs. S12-S24). In Supplementary Movie S7, a sketch of the operating principle about high-content HTFC is shown for the assessment cycle of Fig. 5(a), in which the segmented 3D RI tomograms of single cells flowing and rotating in suspension along a microfluidic channel are characterized by the proposed quantitative biomarkers.

### Investigating modifiers and therapeutic interventions to normalize the lysosomal compartment in NPC cells

We employ the assessed single-cell analysis based on the label-free tomograms to test the effects of two possible therapeutic strategies to rescue cells from the lysosomal mislocalization in NPC disease (experiments D and E in Table S1), as sketched in Fig. 5(b). In experiment D (top row in Fig. 5(b)), we treated HeLa NPC1 KO cells with cyclodextrin, a drug currently under clinical investigation for the NPC treatment [38]. The high-content confocal analysis in the top row of Fig. S6(a) shows that cyclodextrin reduces cholesterol storage in the lysosomal compartment of NPC1 KO cells, although we observed a weak amelioration in the perinuclear accumulation of lysosomes (bottom row in Fig. S6(b)). Instead, in experiment E (bottom row in Fig. 5(b)), to investigate whether we can rescue lysosomal aggregation in the perinuclear area, we used siRNAs to deplete SPAG9, a protein that can drive retrograde lysosomal trafficking and might modify lysosomal positioning [39]. Indeed, we observed that the depletion of SPAG9 reduces the aggregation of lysosomes in HeLa NPC1 KO cells. Most importantly, high-content confocal analysis showed that the repositioning of lysosomes in HeLa NPC1 KO cells lacking SPAG9 was sufficient to significantly reduce cholesterol accumulation, unveiling a potential pathological role of SPAG9 in the NPC disease, thus opening a new therapeutic approach for its treatment (Fig. S7).

We characterized the HeLa NPC1 KO + Cyclodextrin cells and the HeLa NPC1 KO + siSPAG9 cells collected in experiments D and E, respectively, by the proposed HTFC feature set in respect to their control populations, i.e. the HeLa NPC1 KO cells in both cases. Again, we compared the quantitative fingerprint of the lysosomal accumulation obtained in experiments D and E to the ground truth made of the entire dataset of HeLa WT cells and HeLa NPC1 KO cells.

Both the cyclodextrin treatment and SPAG9 gene silencing aim to partially restore the WT condition, thus the lysosomal accumulation should redistribute uniformly around the nucleus, as shown by tomograms in Fig. 5(b). This is confirmed by the NLNSA parameter in Fig. 6(a) and the LCPSSE_Az-El_ parameter in Fig. 6(d), which quantify the degree of uniform spatial distribution about the LVC compartment around the nucleus. However, siSPAG9 has a greater effect than cyclodextrin in absolute terms. Even in terms of HLVR, the PV is almost null in the cyclodextrin treatment of experiment D (Fig. 6(b)). Furthermore, according to the NNCCD parameter in Fig. 6(c), the SPAG9 gene silencing seems to have a greater ability also in restoring the nuclear position and shape typical of the WT-like condition.

A discussion about the other proposed morphometric biomarkers is reported in the Supplementary Information (Section S2 and Figs. S12-S24). In Supplementary Movie S8, a sketch of the operating principle about high-content HTFC is shown for the drug testing cycle of Fig. 5(b), in which the segmented 3D RI tomograms of single cells flowing and rotating in suspension along a microfluidic channel are characterized by the proposed quantitative biomarkers.

## Discussion and Conclusions

Label-free and quantitative HTFC enables reliable imaging of cells in suspension, offering novel insights into intracellular organization in 3D. To study the lysosomal compartment, we defined ad hoc 3D morphometric biomarkers that characterize lysosomes-nucleus, lysosomes-cell, and nucleus-cell arrangements in health (WT) and disease (NPC1) models. Each biomarker comprises specific features computed in cartesian or spherical coordinates. Ranking these features revealed that the lysosomes-nucleus biomarker best captures lysosomal alterations in NPC1 disease, followed by lysosomes-cell and nucleus-cell biomarkers. Notably, spherical coordinates proved more discriminative than cartesian ones, highlighting the advantage of analyzing suspended cells in a comprehensive 3D space. Enabled by flow cytometry modality [40], this HTFC approach allows advanced single-cell statistical analysis of lysosomal organization, with significant implications for lysosomal biology, LSD diagnosis, and therapeutic development.

In this work, we illustrated proof of concept of such promising High-Content HTFC. To validate our strategy, by keeping the WT vs. NPC1 KO distributions as ground truth, we obtained that the U18666A treatment has a strong effectiveness over the WT cells in emulating the NPC1 disease as well as the gene overexpression has a strong effectiveness over the NPC1 KO cells in partially restoring the WT phenotype. We also tested two possible therapeutic treatments that aim at rescuing the NPC1 phenotype. We observed that SPAG9 siRNA-mediated relocation of lysosomes was more effective than cyclodextrin treatment. Also, cyclodextrin has a quasi-null effect in recovering the WT nuclear position (Fig. 6(c), Fig. S23, and Fig. S24), and a negative effect in recovering the WT nuclear shape (Fig. S16 and Fig. S22). To better quantify and compare the effectiveness of the several treatments, we define the treatment’s effectiveness (TE) by exploiting the t-distributed stochastic neighbor embedding (t-SNE) algorithm [41]. In fact, after the z-score standardization of each of the features over the entire collected dataset reported in Table S1, we perform the t-SNE to reduce the dimensionality of the dataset. Given a certain treated population, we define the TE as the Euclidean distance in the t-SNE space between the centroid of its cluster and the centroid of the corresponding control population. In Fig. 7(a), we show the t-SNE clusters of the two ground truth populations, i.e. WT cells and NPC1 KO cells, which have a Euclidean distance of 24.6. Based on the feature analysis discussed above, the effectiveness of the U18666A treatment (TE=19.8) is higher than the gene overexpression (TE=11.3), most likely due to the transient overexpression, and closer to the ground truth (TE=24.6). More important, based on the feature analysis discussed above, the effectiveness of the siSPAG9 treatment (TE=9.9) is higher than the cyclodextrin treatment (TE=5.5) in reverting perinuclear localization of lysosomes, but lower than the gene overexpression (TE=11.3).

**Fig. 7.**
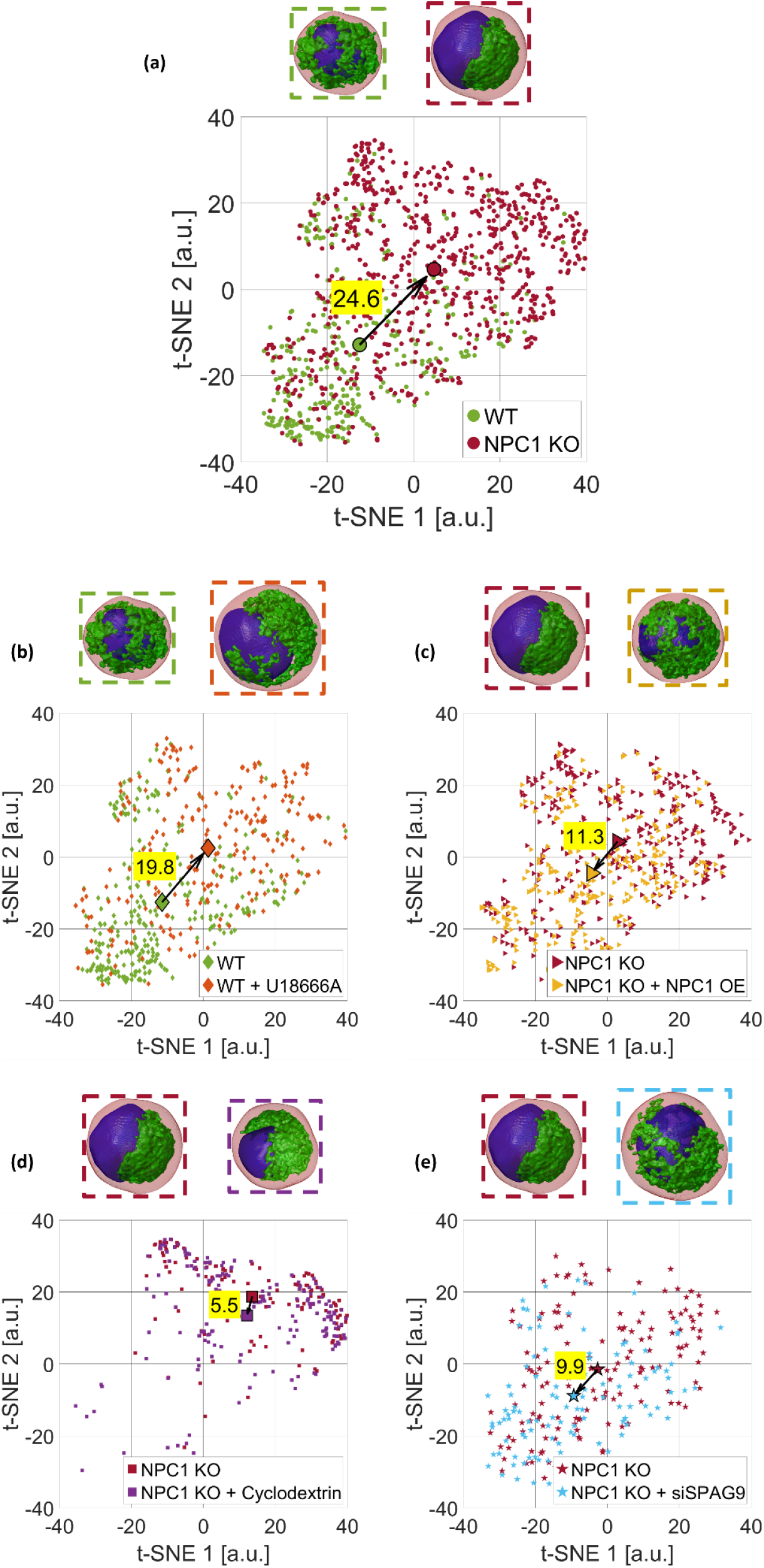
t-SNE visualization of the HTFC feature set used to characterize the label-free 3D RI tomograms of several HeLa cell lines before and after specific biological treatments, thanks to the segmentation of the nucleus (blue) and LVC (green) inside the cell (red). For each experiment, examples of segmented 3D RI tomograms are displayed at the top, the cluster centroid is highlighted, and the Euclidean distance between clusters (i.e., the TE) is reported. **(a)** Ground truth experiment. WT dots are in the bottom-left part of the t-SNE space, while NPC1 KO dots are in the top-right part of the t-SNE space, thus resulting in a TE of 24.6. **(b)** Experiment B. As expected by the scheme in Fig. 1(a), WT + U18666A cells are about in the same position as NPC1 KO cells in (a). **(c)** Experiment C. As expected by the scheme in Fig. 1(a), NPC1 KO + NPC1 OE cells are about in the same position as WT cells in (a). **(d)** Experiment D. As expected by the scheme in Fig. 1(b), NPC1 KO + Cyclodextrin cells are about in the same position as WT cells in (a). **(e)** Experiment E. As expected by the scheme in Fig. 1(b), NPC1 KO + siSPAG9 cells are about in the same position as WT cells in (a).

In summary, we validated single-cell multiparametric biomarkers measured through HTFC for detecting lysosomal and nuclear positioning changes in health and disease conditions by demonstrating their ability to measure alterations induced by a set of pharmacological and genetic manipulations. For the first time in a label-free, single-cell context, we observed that modulating lysosomal positioning directly affects two key LSD hallmarks, i.e. lysosomal swelling and pathological cholesterol accumulation, particularly in NPC1. These findings establish High-Content HTFC as a promising tool for diagnosing lysosomal abnormalities through minimally invasive methods like blood sampling from patients suspected of having LSDs. Additionally, it could assess therapeutic efficacy by monitoring lysosomal recovery. Future studies will expand its application to various LSDs, linking alterations in the lysosomal morphology to specific lysosomal storage pathologies.

## Online Methods

### Sample preparation

HeLa WT and HeLa NPC1 KO cells were cultured in Dulbecco’s Modified Eagle Medium (DMEM) supplemented with 10% fetal bovine serum (FBS, Euroclone), 1% L-glutamine, and 1% penicillin/streptomycin.

In experiments B and D, HeLa WT and HeLa NPC1 KO cells were treated with 1 μM U18666A and 1 mM β-cyclodextrin for 16 hours and 24 hours, respectively.

In experiment C, HeLa NPC1 KO cells were transfected with plasmids NPC1_OHu27327 (GenScript) using Lipofectamine Transfection Reagent (Thermo Fisher Scientific) according to the manufacturer’s instructions. Transfection was performed for 48 hours.

In experiment E, HeLa NPC1 KO cells were transfected with small interfering RNAs (siRNAs) targeting SPAG9/JIP4. Three specific siRNA sequences were used:

- siRNA #1: 5’-GAGUAGUUUAGAUAAGUUATT-3’
- siRNA #2: 5’-GGAAUUAAGUCAACCACGUTT-3’
- siRNA #3: 5’-GGAUCUGACGGGUGACAAATT-3’

Cells were transfected with 40 nM siRNAs using Lipofectamine RNAiMAX reagent (Thermo Fisher Scientific) following the manufacturer’s protocol. Transfection was carried out for 72 hours.

For HTFC experiments, HeLa WT and HeLa NPC1 KO cells were harvested by incubating them for 5 minutes with 0.05% trypsin-EDTA solution (Sigma-Aldrich). Following detachment, cells were resuspended in complete culture medium to a final concentration of 2×10^5^ and then injected into the microfluidic channel.

### High-content confocal imaging

High-content images were captured using the OPERA High Content Imaging System (PerkinElmer). To assess cholesterol accumulation, cells were fixed with 4% paraformaldehyde (PFA) for 10 minutes and washed with phosphate-buffered saline. Cells were permeabilized with blocking buffer for 40 minutes and then stained with 50 μg/mL Filipin (SAE0087, Sigma-Aldrich) for 1 hour at room temperature. Nuclei were counterstained with DRAQ5 (1:5000 dilution; Thermo Fisher Scientific, cat. 62254) or DAPI (1:8000 dilution; Hoechst 33342, Thermo Fisher Scientific, cat.62249) for 10 minutes.

For immunofluorescence, cells were fixed with 4% PFA for 10 minutes, permeabilized with saponin-containing blocking buffer, and incubated with the following primary antibodies:

- LAMP1 (Santa Cruz Biotechnology, cat. sc-20011; 1:400 dilution)
- NPC1 (Abcam, cat. ab134113; 1:200 dilution)

Primary antibody incubation was performed for 1 hour at room temperature, followed by incubation with Alexa Fluor-conjugated secondary antibodies (Thermo Fisher Scientific, AlexaFluor 488 A21202, AlexaFluor 568 A10042; 1:400 dilution) for 45 minutes. Nuclei were counterstained with DRAQ5.

For the quantification of lysosome distribution, high-content confocal images of LAMP1 staining in HeLa WT and HeLa NPC1 KO cells were analyzed by Sima software. The cytoplasm was divided into two defined regions: perinuclear and peripheral. Mean fluorescence intensity of LAMP1 staining was measured separately in both areas. The perinuclear index was calculated as the ratio of the mean intensity in the perinuclear region to that in the peripheral region (perinuclear/peripheral).

### HTFC experimental system

For recording holographic videos, we implemented a Mach-Zehnder interferometer based on an off-axis configuration (Fig. 8(a)) [42]. We use a solid state continuous wave laser as laser source (Laser Quantum Torus 532; wavelength = 532 nm; output power = 750 mW) and a polarizing beam splitter (PBS) to separate the generated light wave into an object and a reference beam. Two half-wave plates (HWPs) are placed in front of and behind the PBS for adjusting the splitting ratio of the two contributions. The object beam passes through the cells that flow within the microfluidic channel (cross section = 200 μm × 1000 μm; length = 58.5 mm; Microfluidic ChipShop). A microscope objective (MO1) with high numerical aperture (Zeiss; ×40; oil immersion; numerical aperture = 1.3) collects the scattered light that is successively sent to a lens (L1) with a focal distance of 150 mm. The reference beam is driven to a second microscope objective (MO2) and a second lens (L2, focal distance = 150 mm). The two wave contributions are recombined by a beam splitter cube (BS) with a non-null angle, according to the definition of the off-axis configuration. A CMOS recording camera (Genie Nano-CXP Camera; 5120 × 5120 pixels, Δx = Δy = 4.5 μm pixel size) acquires the resulting interference fringe pattern, i.e. the hologram (Fig. 8(b)). Cells are injected into the microfluidic channel thanks to an automatic syringe pump (CETONI Syringe Pump neMESYS 290N). The channel form factor ensures a laminar flow such that the cells rotate, if they do not flow in the center of the channel, leveraging the velocity gradient due to the parabolic velocity profile. Therefore, cells undergo roto-translation within the channel, enabling the acquisition of a holographic video sequence of the same cell from different views. In particular, according to the reference system in Figs. 8(a,b), cells flow along the y-axis, rotate around the x-axis, and are acquired along the optical z-axis.

**Fig. 8.**
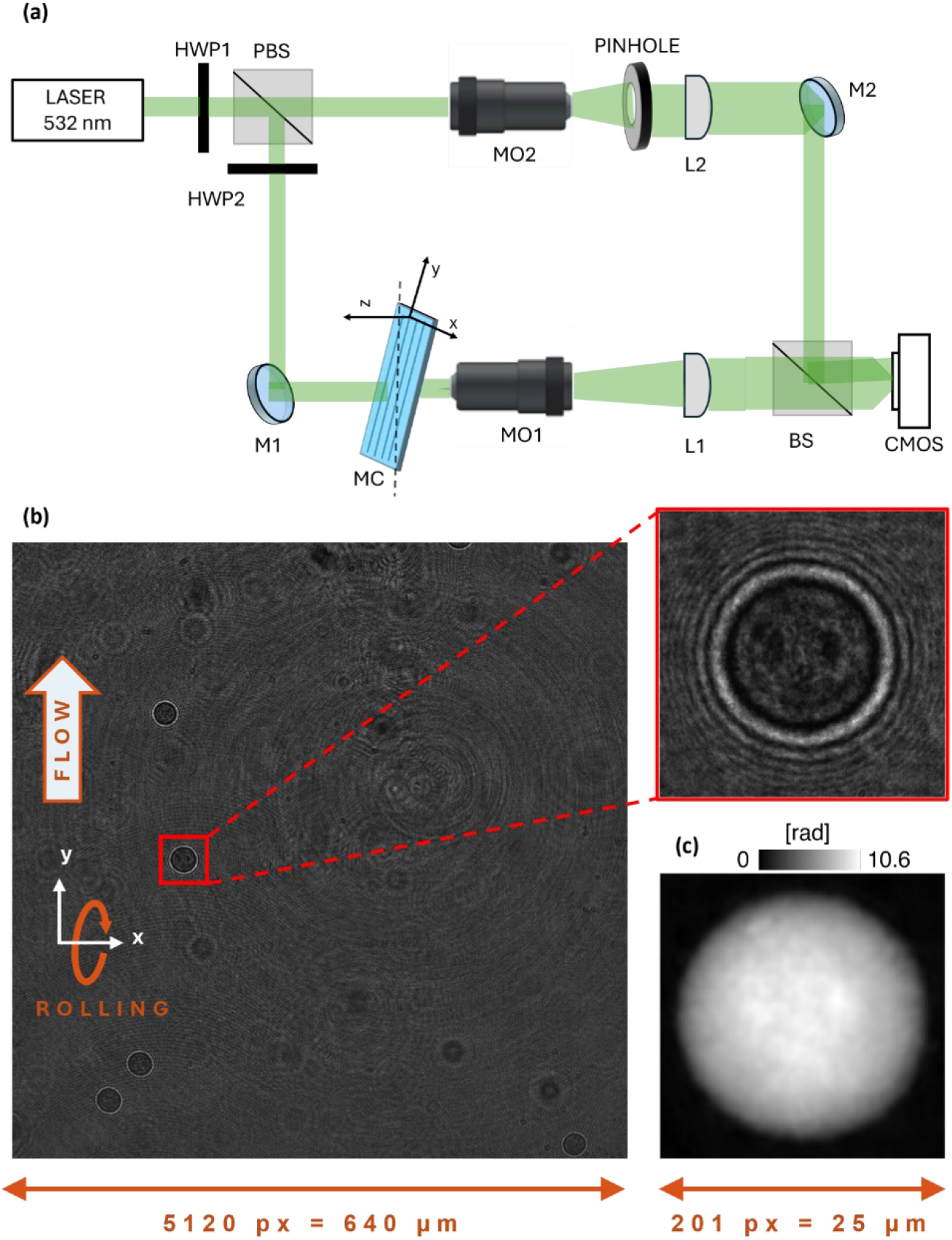
HTFC experiment and numerical reconstruction. **(a)** Opto-fluidic recording system based on a digital holographic microscope. HWP – Half-Wave Plate; PBS – Polarizing Beam Splitter; M – Mirror; L – Lens; MO – Microscope Objective; MC – Microfluidic Channel; BS – Beam Splitter; CMOS – Camera. **(b)** Full hologram recorded during in-flow experiments, with highlighted in red the 384×384 ROI used for detecting and tracking the cell. Cells flow along the y-axis and rotate around the x-axis. **(c)** QPM numerically reconstructed from the cell’s hologram highlighted in (b).

### HTFC numerical reconstruction

By exploiting the inherent contrast between cells and their background, the flowing and rotating single cells are detected and tracked within each frame of the recorded holographic video sequence by using a region of interest (ROI) of sizes 384×384 pixels, as highlighted in red in Fig. 8(b). Then, for each cell, after identifying their centroids during roto-translation, a sequence of 1024×1024 sub-holograms is cropped from the recorded holographic video sequence to allow reconstructing their corresponding Quantitative Phase Maps (QPMs) [21,22]. Thus, on each sub-hologram, we perform the same numerical reconstruction pipeline. The first step is the hologram demodulation, applying the Fourier spectrum filtering for selecting and centering the real diffraction order [42]. Then, the demodulated hologram is numerically propagated along the z-axis through the Angular Spectrum method [42] in order to obtain the proper in-focus distance by minimization of the Tamura coefficient [43]. By numerically propagating the demodulated hologram to the evaluated distance, we obtain the in-focus complex amplitude. The retrieved phase-contrast image (i.e. the calculated argument of the complex amplitude) is optimized by removing the residual optical aberrations by using a reference hologram [44]. Finally, we employ the two-dimensional windowed Fourier transform filtering [45] and the PUMA algorithm [46] for denoising and unwrapping steps, respectively, thus obtaining the QPM (Fig. 8(c)). Each QPM is centered with respect to the cell’s centroid to avoid motion artifacts during the tomographic reconstruction. Then, by recognizing phase similarities among all the QPMs of the same flowing and rolling cell, the unknown viewing angles are estimated from the tracking positions [47]. Finally, the centered QPMs and the corresponding viewing angles of a flowing and rotating cell are given in input to the Filtered Back Projection algorithm [48] to compute its 3D RI tomogram (Figs. 2(a,f)).

### Segmentation of lysosomal aggregates in 3D RI tomograms

By exploiting the high RI values of lysosomal compartment [18,33,34], we start lysosomal segmentation by establishing a rough estimate of the lysosomal volume by setting as threshold the 0.80-quantile computed from the RI statistical distribution of the overall 3D cell tomogram (Figs. 2(b,g)). Then, we implement the CSSI algorithm to segment the nucleus inside the same cell [20,21]. The CSSI algorithm is based on the iterative execution of statistical hypothesis tests to compare several groups of intracellular RI voxels with respect to a reference group, supposed to belong to the organelle to be segmented. Usually, the central voxels of the cell belong to the nucleus [20]. Therefore, we divide the *L*_*x*_ × *L*_*y*_ × *L*_*z*_ array containing the 3D RI tomograms into non-overlapping cubes with *ε* side (i.e., they are made of *ε*^3^ RI values), and we choose the reference group as the *ε*-cube closest to the cell center without having voxels belonging to the rough lysosomal volume, as highlighted in yellow in Figs. 2(b,g). In this way, the boundaries between the nuclear compartment and the lysosomal compartment can be better defined since there is no statistical ambiguity between the two corresponding RI statistical distributions. Finally, after the CSSI segmentation of the nuclear compartment (Figs. 2(c,h)), the rough lysosomal volume is refined by deleting all voxels within the nuclear boundaries (Figs. 2(d,i)). In fact, because of the partial overlapping between the RI statistical distributions of different organelles [28], it could happen that the same RI threshold used for segmenting the lysosomal compartment is able to collect also the highest RI voxels inside the nucleus. The segmented nuclear and lysosomal compartments can be observed together in Figs. 2(e,j) for the HeLa WT cell and the HeLa NPC1 KO cell, respectively.

## Supplementary Material

*Supplementary Movie S1*. Steps of the algorithm for the multi-specific segmentation of nucleus and lysosomes in a suspended HeLa WT cell recorded and reconstructed by Holo-Tomographic Flow Cytometry.

*Supplementary Movie S2*. Steps of the algorithm for the multi-specific segmentation of nucleus and lysosomes in a suspended HeLa NPC1 KO cell recorded and reconstructed by Holo-Tomographic Flow Cytometry.

*Supplementary Movie S3*. Visual comparison between the multi-specific 3D images from fluorescence High-Content Confocal Microscopy and label-free High-Content Holo-Tomographic Flow Cytometry about an adherent HeLa WT cell and a suspended HeLa WT cell, respectively.

*Supplementary Movie S4*. Visual comparison between the multi-specific 3D images from fluorescence High-Content Confocal Microscopy and label-free High-Content Holo-Tomographic Flow Cytometry about an adherent HeLa NPC1 KO cell and a suspended HeLa NPC1 KO cell, respectively.

*Supplementary Movie S5*. Visualization of the half-nuclear lysosomes volume ratio (HLVR) feature measuring the spatial distribution of lysosomes on two sides of the nucleus divided in half by a cutting plane in some suspended HeLa WT cells and HeLa NPC1 KO cells imaged by High-Content Holo-Tomographic Flow Cytometry.

*Supplementary Movie S6*. Visualization of the normalized lysosomes-nucleus solid angle (NLNSA) feature measuring the solid angle that subtends lysosomes from the nucleus centroid in some suspended HeLa WT cells and HeLa NPC1 KO cells imaged by High-Content Holo-Tomographic Flow Cytometry.

*Supplementary Movie S7*. Sketch of the operating principle about High-Content Holo-Tomographic Flow Cytometry for the assessment cycle of lysosomal accumulation in which the segmented 3D RI tomograms of single HeLa cells flowing and rotating in suspension along a microfluidic channel are characterized by the lysosomes-nucleus biomarker.

*Supplementary Movie S8*. Sketch of the operating principle about High-Content Holo-Tomographic Flow Cytometry for the drug testing cycle of lysosomal accumulation in which the segmented 3D RI tomograms of single HeLa cells flowing and rotating in suspension along a microfluidic channel are characterized by the lysosomes-nucleus biomarker.

## Acknowledgements

We thank the High Content Screening Facility at TIGEM. We acknowledge financial support from the Italian Telethon Foundation.

This work was partially supported by project “FIGHT-LSDs” – Flow-cytometry ImaGing by Holographic Tomography to advance study, diagnosis and treatment of Lysosomal Storage Diseases – funded by *Accordo per la Coesione della Regione Campania. Fondo di Rotazione ex L. 183/1987* (project n. 48).

This work was partially supported by project “CITOM” – Programma AMICO 2, CNR – UVR - within the PoC 2022 – PNRR funded by the Italian Ministry of Business and Made in Italy – UIBM in the framework of Next Generation EU.

## Author contributions

Diego Luis Medina and Pietro Ferraro conceptualized the idea behind the study. Lisa Miccio and Pasquale Memmolo designed the holographic experiments and the computational strategies, respectively. Michela Schiavo implemented biological treatments and conducted cell cultures. Giusy Giugliano and Sandro Montefusco performed holographic and confocal experiments, respectively. Daniele Pirone carried out data analyses and biomarkers’ computation. Daniele Pirone, Michela Schiavo, Pasquale Memmolo, Diego Luis Medina, and Pietro Ferraro wrote the original draft. Pasquale Memmolo, Diego Luis Medina, and Pietro Ferraro supervised the study. Diego Luis Medina and Pietro Ferraro managed the acquisition of funding. All authors read and approved the final manuscript.

## Competing interests

The authors declare no completing interests.

## Notes

### Competing Interest Statement

The authors have declared no competing interest.

