## Supplementary Information for "Quantitative profiling of lysosomal accumulation through label-free biomarkers via High-Content Holo-Tomographic Flow Cytometry"

### S1. Set of quantitative 3D morphometric features in HTFC

The HTFC working principle allows to record and reconstruct the 3D RI tomograms of label-free cells flowing and rotating in suspension along a microfluidic channel. Thus, a comprehensive 3D morphometric analysis can be carried out to quantitatively characterize the real arrangement of lysosomes inside the whole cellular volume in respect to the nucleus and the cytoplasm, in order to extract a fingerprint that is as distinctive as possible of the NPC disease. At this aim, here we propose an ad hoc feature set made of 17 morphometric parameters computed in the entire 3D space. The proposed morphometric parameters are listed in Table S2 along with the corresponding acronyms and value ranges, and their categorization into several classes is shown Fig. S8.

The *half-nuclear lysosomes volume ratio (HLVR)* measures how uniformly distributed the lysosomes are around the nucleus. At this aim, the quasi-spherical nucleus is divided into two parts by a cutting plane passing through the nucleus centroid and orthogonal to the segment joining the nucleus centroid to the LVC centroid, as sketched in the example of Figs. 4(a,d), containing the same cells of Figs. 2(e,j), respectively. Let $V_{LVC}^{+}$ be the LVC volume in the half-space containing the LVC centroid, let $V_{LVC}^{-}$ be the LVC volume in the other half-space, and let

$$\begin{aligned} HLVR^{+} &= \frac{V_{LVC}^{+}}{V_{LVC}^{+}+V_{LVC}^{-}} = 1 - HLVR^{-} \\ HLVR^{-} &= \frac{V_{LVC}^{-}}{V_{LVC}^{+}+V_{LVC}^{-}} = 1 - HLVR^{+} \end{aligned} . \quad \text{(S1)}$$

We arbitrarily define the $HLVR$ as the $HLVR^{+}$, which can take values from 0.5 (lysosomes are uniformly distributed around the nucleus) to 1 (lysosomes are accumulated on a single side of the nucleus). In the example of Figs. 4(a,d), $HLVR = 0.67$ in the case of the HeLa WT cell (Fig. 4(a)) and $HLVR = 1$ in the case of the HeLa NPC1 KO cell (Fig. 4(d)). Indeed, it can be seen that, as expected, the HeLa NPC1 KO cell in Fig. 4(d) has a highly asymmetric lysosomal accumulation, which is well quantified by the HLVR with respect to the much more uniform perinuclear distribution of lysosomes in the HeLa WT cell of Fig. 4(a). To further verify this, we computed the HLVR parameter over the entire dataset of 358 HeLa WT cells and 766 HeLa NPC1 KO cells. In the insets of Figs. 4(a,d), we show the pie chart with the average $HLVR^-$ and $HLVR^+$ values over the whole population of HeLa WT cells and HeLa NPC1 KO cells, respectively. As expected, in the disease condition of HeLa NPC1 KO cells (inset in Fig. 4(d)), the sizes of the two slices of pie are much more unbalanced than the healthy condition of HeLa WT cells (inset in Fig. 4(a)), thus quantifying an accumulation of the lysosomal compartment on one side of the nucleus.

Another parameter here introduced to measure the lysosomal accumulation with respect to the nucleus is the *normalized lysosomes-nucleus solid angle (NLNSA)*. After centering the coordinate reference system in the nucleus centroid, we compute the solid angle that subtends the LVC, as shown in the example of Figs. 4(b,f) containing the same cells of Figs. 2(e,j), respectively. Then, we normalize the solid angle to $4\pi$ so that the NLNSA biomarker can take values in the $[0,1]$ range. In particular, high values are related to uniform distribution of lysosomes around the nucleus, while low values are related to a lysosomal accumulation around the nucleus. For example, $NLNSA = 1$ in the HeLa WT cell of Fig. 4(b), while $NLNSA = 0.45$ in the HeLa NPC1 KO cell of Fig. 4(e).

A further quantitative feature we define is the *normalized lysosomes-nucleus centroids distance (NLNCD)*, that measures the distance between the nucleus centroid and the LVC centroid (see the white line in Figs. 4(a,d)), normalized to the cell's equivalent radius, that is the radius of a sphere having the same volume as the cell.

The *normalized lysosomes-nuclear membrane distance (NLNMD)* measures instead the average distance between each voxel of the LVC compartment and its closest voxel belonging to the nuclear membrane, normalized to the cell's equivalent radius. The *lysosomes-cell surface ratio (LCSR)* measures the surface area of the LVC compartment, normalized to the cell's surface area. The *lysosomal sphericity (LS)* measures the sphericity of the LVC compartment, which is defined as the ratio between the surface area of a sphere having the same volume as the object (i.e., the LVC compartment in this case) and the object's surface area. Sphericity is usually employed to quantify the roundness of an object. Instead, due to the not rounded geometry of the LVC compartment, here we employ the sphericity index to relate the LVC volume to the LVC surface area, thus obtaining a parameter normalized in the $[0,1]$ range able to measure the degree of volumetric compactness of the lysosomal spatial distribution. Instead, the *normalized nuclear sphericity (NNS)* measures the sphericity of the nucleus, and, in this case, it is used to quantify the roundness of this intracellular compartment. In particular, as the nucleus has a quasi-spherical shape, its sphericity is in the $[0.9,1]$ range in most cases, thus the NNS is obtained after subtracting $0.9$ to the nuclear sphericity and after dividing the result by $0.1$, in order to increase the dynamic of this parameter. Finally, the *normalized nucleus-cell centroids distance (NNCCD)* measures the distance between the nucleus centroid and the cell centroid, normalized to the cell's equivalent radius.

Besides, other three feature sets can be defined by converting the cartesian coordinates of the intracellular components into the corresponding spherical coordinates and by considering the azimuth and the elevation angles, as shown in the illustrative example of Figs. 4(c,f).

The first spherical feature set is obtained by centering the spherical coordinate system into the nucleus centroid, as shown in the HeLa WT cell and HeLa NPC1 KO cell of Figs. S9(a,e), respectively, that are the same cells of Figs. 2(e,j) and Fig. 4. This first spherical feature set measures how lysosomes are uniformly distributed around the nucleus. Therefore, a perinuclear volume is computed by subtracting the nuclear volume to its isotropic morphological dilation [S1] based on a spherical structuring element with radius $R$ (in this work, $R = 10\, pixels$), as displayed in yellow in Figs. S9(a,e). For both the perinucleus and the LVC, we calculate the azimuth and the elevation coordinates of their voxels in the chosen reference system. We compare the histograms of the azimuth coordinates of the perinucleus and the LVC in Figs. S9(b,f), while the histograms of the elevation coordinates are reported in Figs. S9(c,g). If the lysosomes are uniformly distributed around the nucleus, the histogram of the LVC azimuth/elevation angles is expected to be similar to the histogram of the perinucleus azimuth/elevation angles. In fact, the LVC and perinucleus histograms are much more similar each other in the WT case of Figs. S9(b,c) than the NPC1 KO case of Figs. S9(f,g). In particular, in both the WT and NPC1 KO cases of Figs. S9(b,f), respectively, due to the quasi-spherical shape of the nucleus, the perinucleus azimuth histogram is quite uniform. Instead, due to the lysosomal accumulation, the LVC azimuth histogram is much peaked in the NPC1 KO case of Fig. S9(f), while it is rather uniform in the WT case of Fig. S9(b). Hence, to quantify this property, we exploit the PSSE, defined as

$$PSSE = 100\frac{\sum(h-h_R)^2}{\sum h_R{}^2}, \qquad \text{(S2)}$$

where $h$ is the histogram of the test object, normalized to its maximum, and $h_R$ is the histogram of the reference object, normalized to its maximum. By using the perinucleus as reference object and the LVC as test object, the *lysosomes-nucleus PSSE of the azimuth coordinate (LNPSSE$_{Az}$)* and the *lysosomes-nucleus PSSE of the elevation coordinate (LNPSSE$_{El}$)* can be computed. As expected, they take much lower values in the WT case of Figs. S9(b,c) (i.e., $LNPSSE_{Az} = 8.81$ % and $LNPSSE_{El} = 3.85$ %, respectively) than the NPC1 KO case of Figs. S9(f,g) (i.e., $LNPSSE_{Az} = 206.92$ % and $LNPSSE_{El} = 31.77$ %, respectively). Furthermore, we also compute the bivariate histogram of the azimuth and elevation angles (Figs. S9(d,h)) in order to perform a joint comparison between the spherical coordinates of the LVC and nuclear compartments. Also in this case, the bivariate histograms of the HeLa WT cell about the perinucleus and the LVC (top and bottom of Fig. S9(d), respectively) are much more similar each other than the corresponding bivariate histograms of the NPC1 KO cell (top and bottom of Fig. S9(h)). This is also confirmed by the *lysosomes-nucleus PSSE of both the azimuth coordinate and elevation coordinates (LNPSSE$_{Az\text{-}El}$)*, which is computed by using the bivariate histograms in the Eq. (S2). Indeed, the LNPSSE$_{Az\text{-}El}$ is much lower in the WT case of Fig. S9(d) (i.e., $LNPSSE_{Az-El} = 44.61$ %) than the NPC1 KO case of Fig. S9(h) (i.e., $LNPSSE_{Az-El} = 355.75$ %).

The second spherical feature set is obtained by centering again the spherical coordinate system into the nucleus centroid, as shown in the HeLa WT cell and HeLa NPC1 KO cell of Figs. S10(a,e), respectively, that are the same cells of Figs. 2(e,j) and Fig. 4. However, in this case, the degree of uniformity of the lysosomal spatial distribution can be evaluated inside the cytoplasm (i.e., the whole cell volume without the nucleus). At this purpose, in the Eq. (S2), we consider the cytoplasm as reference object and the LVC as test object, thus computing the *lysosomes-cytoplasm PSSE of the azimuth coordinate (LCPSSE$_{Az}$)*, the *lysosomes-cytoplasm PSSE of the elevation coordinate (LCPSSE$_{El}$)*, and the *lysosomes-cytoplasm PSSE of both the azimuth coordinate and elevation coordinates (LCPSSE$_{Az\text{-}El}$)*, which are shown in Fig. S10. In this lysosomes-cytoplasm feature set, the same considerations made for the lysosomes-nucleus feature set apply, and thus the LCPSSE values are higher in the NPC1 KO case than the WT case because of the lysosomal accumulation.

Finally, the third spherical feature set is obtained by centering the spherical coordinate system into the cell centroid, as shown in the HeLa WT cell and HeLa NPC1 KO cell of Figs. S11(a,e), respectively, that are the same cells of Figs. 2(e,j) and Fig. 4. In fact, in this case the LVC is neglected as the goal is to measure the spatial positioning of the nucleus inside the cell by means of some parameters different from the simpler nucleus-cell centroid distance. Therefore, in the Eq. (S2), we consider the whole cell as reference object and the nucleus as test object, thus computing the *nucleus-cell PSSE of the azimuth coordinate (NCPSSE$_{Az}$)*, the *nucleus-cell PSSE of the elevation coordinate (NCPSSE$_{El}$)*, and the *nucleus-cell PSSE of both the azimuth coordinate and elevation coordinates (NCPSSE$_{Az\text{-}El}$)*, which are shown in Fig. S11. In particular, as the cell has a quasi-spherical shape, its azimuth coordinates have a uniform distribution (Figs. S11(b,f)) and its elevation coordinates have a gaussian distribution (Figs. S11(c,g)) in both the WT and NPC1 KO cell. Instead, the nucleus is concentric with the cell in the WT case (Fig. S11(a)), while it is horizontally decentralized in the NPC1 KO case (Fig. S11(e)). As a consequence, the nucleus elevation coordinates have a distribution similar to the cell ones in both the WT and the NPC1 KO cases (Figs. S11(c,g), respectively). Instead, the nucleus azimuth coordinates have a distribution similar to the cell ones only in the WT case (Fig. S11(b)), while it diverges in the NPC1 KO case (Fig. S11(f)). For this reason, the NCPSSE$_{El}$ has low values in both the WT and the NPC1 KO cases, while the NCPSSE$_{Az}$ and the NCPSSE$_{Az\text{-}El}$ takes

values much higher in the NPC1 KO case than the WT one. Hence, high values of the NCPSSE indicate a decentralization of the nucleus with respect to the whole cell.
To rank morphometric features according to their ability in discerning the NPC healthy or diseased single-cell state, we calculated a specific score, i.e. the Fisher's discriminant ratio (FDR) [35], as

$$FDR = \frac{(\mu_{wt}-\mu_{ko})^2}{\sigma_{wt}^2+\sigma_{ko}^2}, \quad \text{(S3)}$$

where $\mu_{wt}$ and $\sigma_{wt}$ are the mean value and standard deviation of the feature of a HeLa WT population, and $\mu_{ko}$ and $\sigma_{ko}$ are the mean value and standard deviation of the feature of the HeLa NPC1 KO population. The 17 proposed features are thus sorted in Fig. 3(a) based on their FDR score.

### S2. Comparative analysis among several HTFC experiments based on quantitative 3D morphometric biomarkers

To quantify changes in a certain feature due to a specific treatment, we measure the percentage variation (PV), defined as

$$PV = 100\frac{Me-\overline{Me}}{\overline{Me}}, \quad \text{(S4)}$$

where $Me$ is the median value of the feature of a certain cell population, and $\overline{Me}$ is the median value of the feature of the corresponding control population.
Let's consider the assessment cycle of Fig. 5(a). As for the NLNCD parameter (Fig. S12), a positive PV is observed in the ground truth experiment passing from WT cells to NPC1 KO cells and in experiment B emulating the NPC condition, while a negative PV is observed in experiment C restoring the WT condition. In fact, as shown in Figs. 4(a,d), a greater lysosomal accumulation on one side of the nucleus, leading to higher HLVR values, also corresponds to a greater distance between the nucleus and the LVC centroids, leading to higher NLNCD values. Instead, if lysosomes are uniformly distributed around the nucleus, the LVC centroid is much closer to the nucleus centroid, resulting in lower NLNCD values. Moreover, as shown in Figs. 4(a,d), a greater lysosomal accumulation also means a greater thickness of the LVC compartment, which reflects in higher values of the NLNMD parameter in the NPC condition, as also reported in Fig. S13. Therefore, the NLNMD values on average increase with the WT + U18666A cells and decrease with the NPC1 KO + NPC1 OE cells in respect to the corresponding control cell populations (Fig. S13). The other two parameters strictly related to the lysosomal accumulation are the LCSR and the LS, measuring the volumetric compactness of the LVC compartment. In fact, lower LCSR values and higher LS values indicate a greater compactness of the LCV compartment. Therefore, NPC1 KO cells and WT + U18666A cells have a lower LCSR value (Fig. S14) and a greater LS value (Fig. S15) than WT cells in the ground truth experiment and experiment B. Instead, in experiment C, the NPC1 KO + NPC1 OE cells have greater values than the control NPC1 KO cells, even if with a much lower PV (Fig. S14), while the LS values remain almost unchanged after the treatment (Fig. S15). Another effect of the lysosomal accumulation observed in the 3D RI tomograms of suspended HeLa cells is the corresponding change of the nuclear shape. As displayed in the tomograms of Figs. 4(a,d), the nucleus appears more spherical and concentric with the cell in the healthy case when lysosomes are uniformly distributed around it. Instead, the nucleus becomes more flattened towards the cellular membrane due to the lysosomal accumulation typical of the NPC condition. Therefore, as displayed in Fig. S16, the NNS is lower in the NPC1 KO case of the ground truth and in the WT + U18666A case of experiment B than the WT population, while it increases in the NPC1 KO + NPC1 OE case of the rescue experiment C. The other features herein proposed to characterize the lysosomal accumulation are those based on the spherical coordinate system. Higher values of the LNPSSE parameter indicate a greater lysosomal accumulation around the nucleus, higher values of the NCPSSE parameter indicate a greater decentralization and deformation of the nucleus with respect to the cell centroid and shape, and the LCPSSE parameter takes both phenomena about lysosomes and nucleus into account. Therefore, the LNPSSE (Fig. S17, Fig. S18, and Fig. S19), LCPSSE (Fig. 6(d), Fig. S20, and Fig. S21), and NCPSSE (Fig. S22, Fig. S23, and Fig. S24) values are higher in the NPC1 KO case of the ground truth and in

the WT + U18666A case of experiment B than the WT population, while they decrease in the NPC1 KO + NPC1 OE case of the rescue experiment C.

Let's consider the drug testing cycle of Fig. 5(b). In both experiments D and E (i.e., cyclodextrin and SPAG9 gene silencing, respectively), the NLNMD parameter has a similar PV passing from the NPC1 KO + Cyclodextrin and NPC1 KO + siSPAG9 to the control WT cells, opposite in sign to the PV passing from WT cells to NPC1 KO cells (Fig. S13). Instead, while maintaining the opposite sign with respect to the ground truth, the PV is much lower in the cyclodextrin treatment of experiment D than the SPAG9 gene silencing of experiment E for the NLNCD (Fig. S12). In terms of volumetric compactness of the LVC compartment, cyclodextrin and siSPAG9 have a similar behavior opposite in sign to the ground truth experiment, as reported in Fig. S14 and Fig. S15 by the LCSR and LS, respectively. In terms of nuclear sphericity measured by the NNS (Fig. S16), while the SPAG9 gene silencing provides a more spherical shape to the nucleus typical of the WT case, the cyclodextrin treatment even makes the nuclear shape less spherical with respect to the control NPC1 KO condition. Moreover, the different effects on nuclear properties can be observed also in the NCPSSE features measured in a spherical coordinate system. In particular, the PV signs are discord between experiments D and E in terms of NCPSSE$_{Az}$ (Fig. S22), as cyclodextrin PV is concord with the ground truth, while it should have an opposite effect as well as siSPAG9. Instead, in terms of NCPSSE$_{El}$ (Fig. S23) and NCPSSE$_{Az\text{-}El}$ (Fig. S24), the PV is opposite in sign to the ground truth in both experiments D and E, but, in absolute terms, siSPAG9 has a greater effect than cyclodextrin. This occurs also with the LNPSSE$_{Az}$ (Fig. S17) and the LCPSSE$_{Az}$ (Fig. S20), while the LNPSSE$_{Az\text{-}El}$ (Fig. S19), the LNPSSE$_{El}$ (Fig. S18), the LCPSSE$_{Az\text{-}El}$ (Fig. 6(d)), and the LCPSSE$_{El}$ (Fig. S21) exhibit more similar PV values of cyclodextrin and siSPAG9 treatments, both opposite in sign to the ground truth.

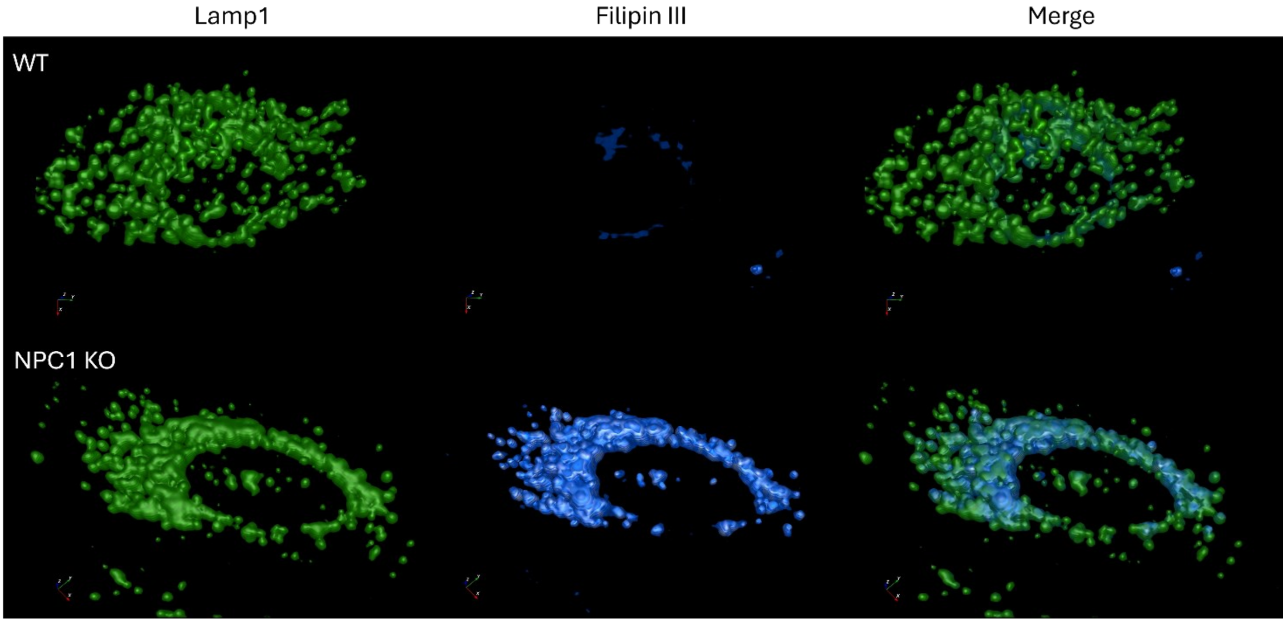


**Fig. S1. 3D representation of a (top row) HeLa WT cell and a (bottom row) HeLa NPC1 KO cell acquired by high-content confocal imaging.** Cells are stained using Lamp1 (green) to detect lysosomes and Filipin III (blue) to detect cholesterol. Images show a clear difference of cholesterol storage inside lysosomes between the two cell lines and a different distribution of lysosomes in the perinuclear area, which is much more uniform in the WT case than the NPC1 KO case.

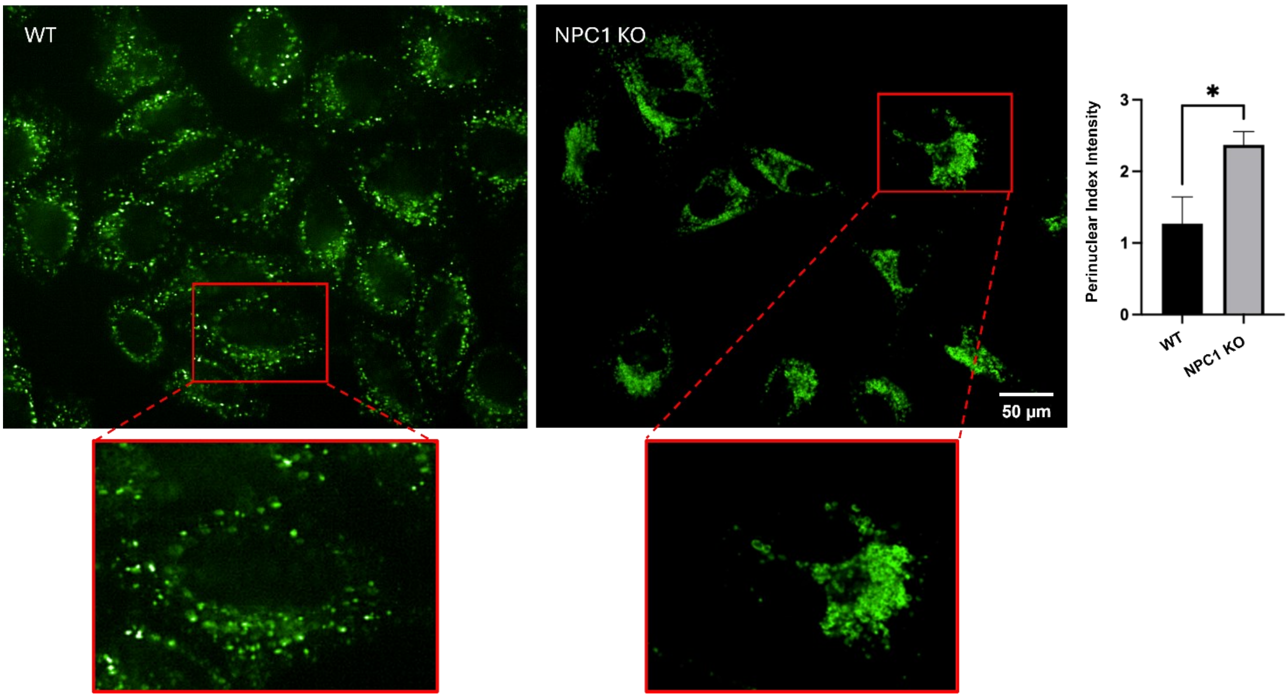


**Fig. S2. Slice representation of the lysosomal compartment stained by Lamp1 (green) in (on the left) HeLa WT cells and (on the right) HeLa NPC1 KO cells acquired by high-content confocal imaging.** Lysosomes in NPC1 KO cells show a perinuclear clustering, in contrast to the more uniform distribution observed in WT cells. For the quantification of the lysosomal distribution inside the cell, the cytoplasm is segmented into two areas (perinuclear and peripheral). The perinuclear index intensity on the right reports the ratio between the perinuclear and peripheral mean intensities related to Lamp1. Higher values mean greater perinuclear accumulation of lysosomes.

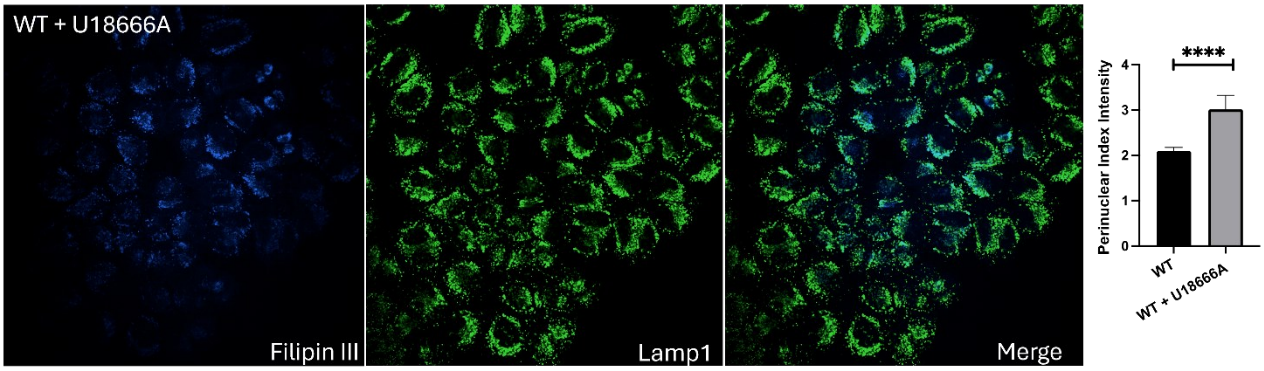


**Fig. S3. HeLa WT cells treated with U18666A compound (namely, WT + U18666A) show a strong accumulation of lysosomes in perinuclear areas within high-content confocal images.** Cells are stained using Lamp1 (green) to detect lysosomes and Filipin III (blue) to detect cholesterol. For the quantification of the lysosomal distribution inside the cell, the cytoplasm is segmented into two areas (perinuclear and peripheral). The perinuclear index intensity on the right reports the ratio between the perinuclear and peripheral mean intensities related to Lamp1. Higher values mean greater perinuclear accumulation of lysosomes.

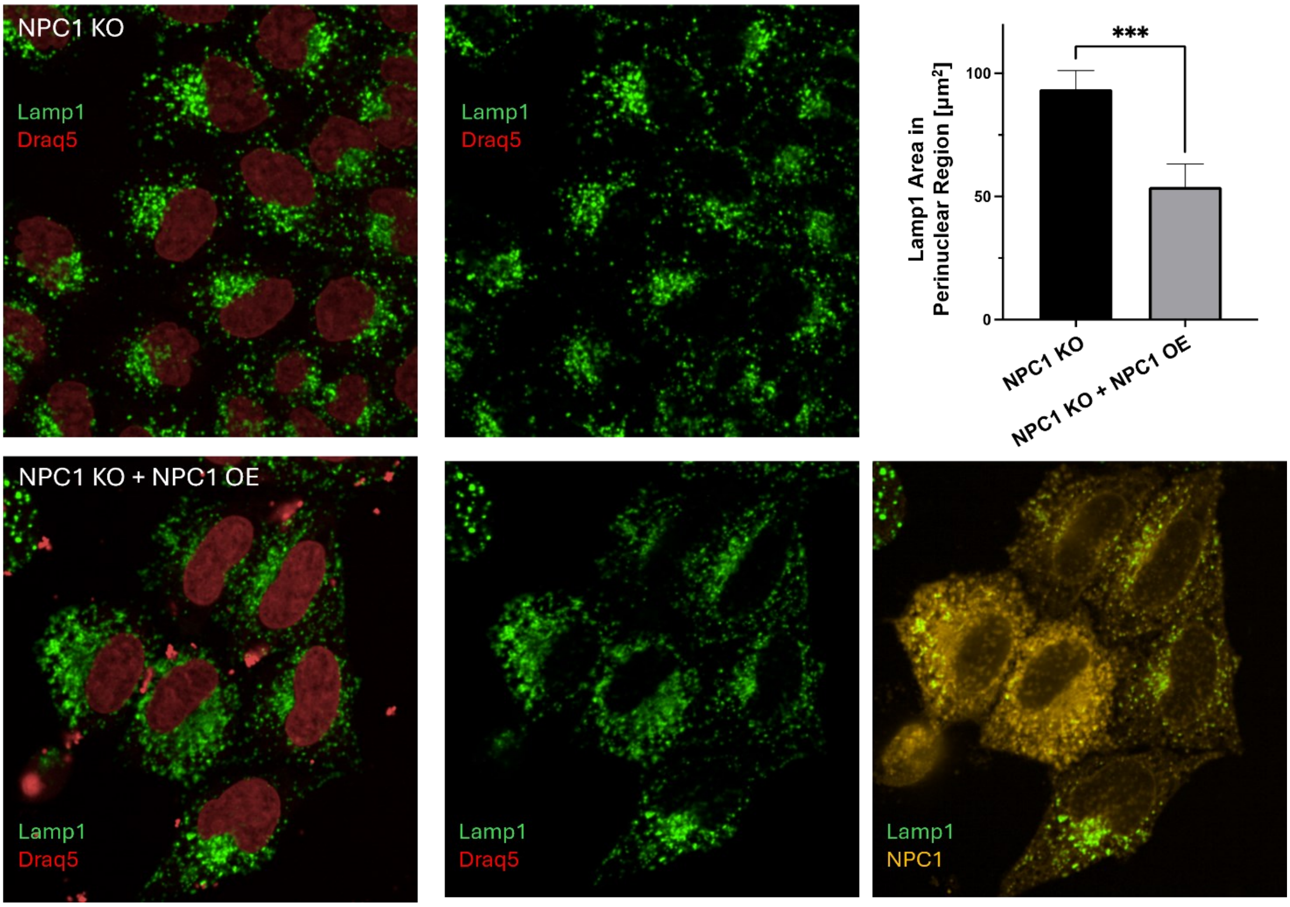


**Fig. S4. Transient overexpression (OE) of the disease-causing NPC1 gene relocates lysosomes to a more uniform distribution in (bottom row) HeLa NPC1 KO cells (namely, NPC1 KO + NPC1 OE) with respect to (top row) control HeLa NPC1 KO cells acquired by high-content confocal imaging.** Cells are stained using Lamp1 (green) to detect lysosomes, Draq5 (red) to detect nuclei, and NPC1 (yellow) to detect the protein. The Lamp1 area in the perinuclear region of transfected and non-transfected cells is reported at the top right, demonstrating a reduction in lysosomal storage in the transfected cells.

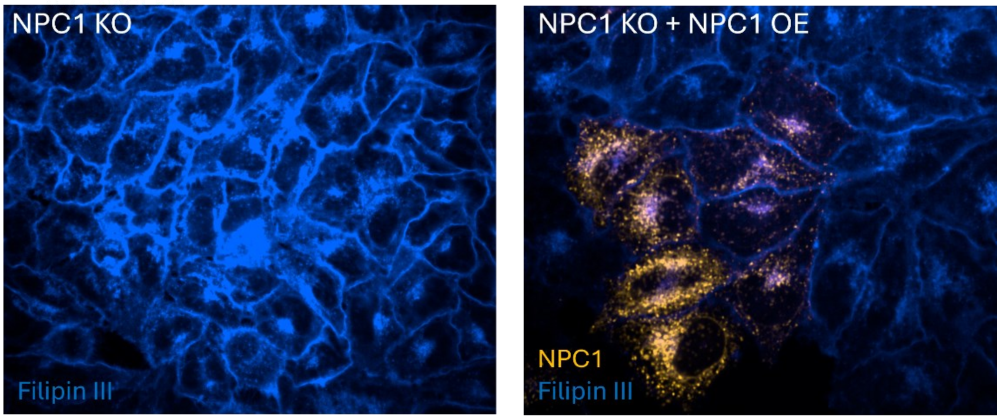


**Fig. S5. Overexpression (OE) of the NPC1 gene reduces Filipin III-stained cholesterol accumulation in (on the right) mutant HeLa cells (namely, NPC1 KO + NPC1 OE) with respect to the (on the left) control HeLa NPC1 KO cells acquired by high-content confocal imaging.** Cells are stained using Filipin III (blue) to detect cholesterol and NPC1 (yellow) to detect the protein.

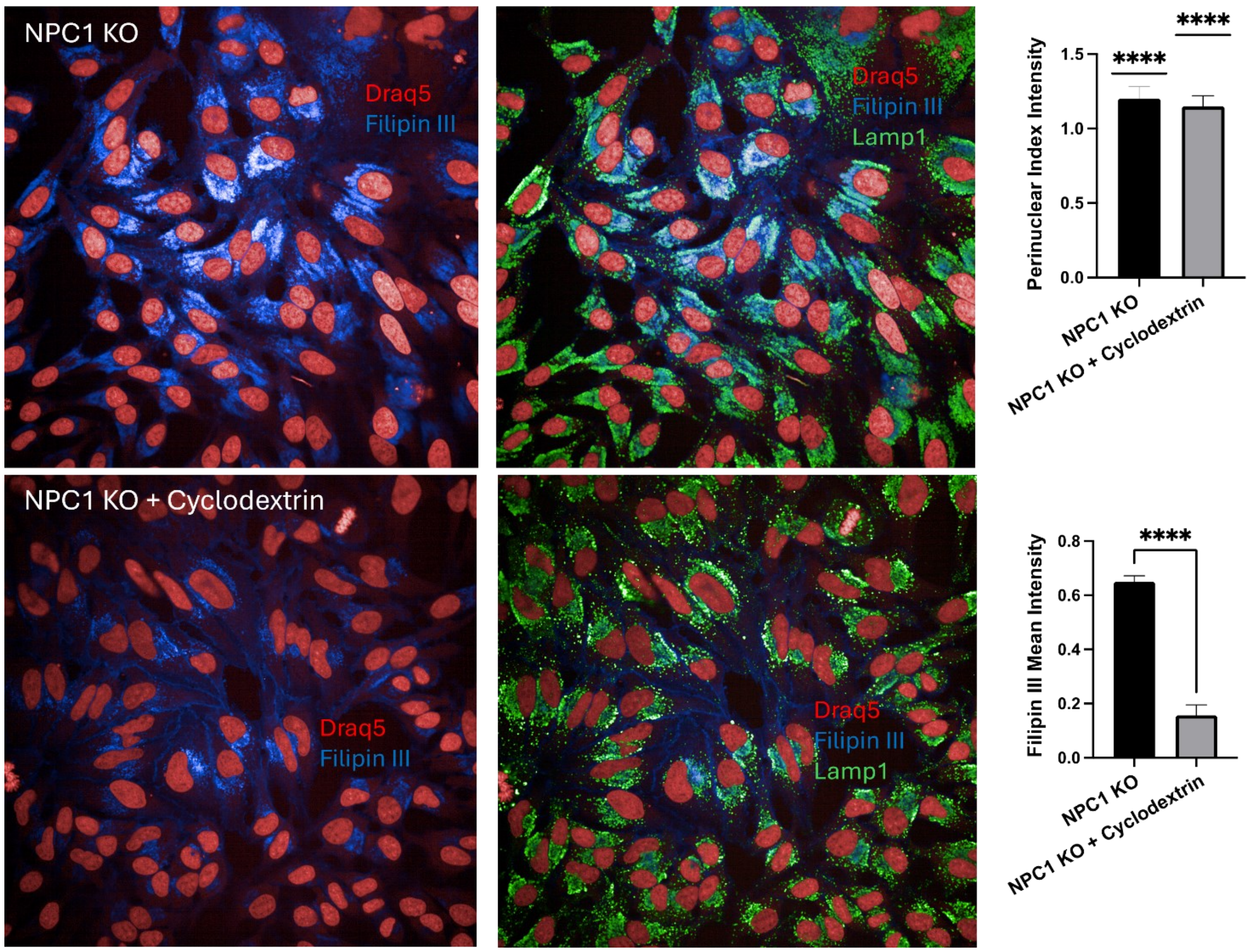


**Fig. S6. Treatment with cyclodextrin reduces cholesterol storage in the lysosomal compartment of (bottom row) mutant HeLa cells (namely, NPC1 KO + Cyclodextrin) with respect to the (top row) control HeLa NPC1 KO cells, as shown in Filipin III assay imaged by high-content confocal imaging.** Cells are stained using Lamp1 (green) to detect lysosomes, Filipin III (blue) to detect cholesterol, and Draq5 (red) to detect nuclei. For the quantification of the lysosomal distribution inside the cell, the cytoplasm is segmented into two areas (perinuclear and peripheral). The perinuclear index intensity at top right reports the ratio between the perinuclear and peripheral mean intensities related to Lamp1. As higher values mean greater perinuclear accumulation of lysosomes, it shows a weak reduction of the lysosomal accumulation in the perinuclear area after cyclodextrin treatment. The mean intensity of Filipin III at bottom right is instead strongly reduced after cyclodextrin treatment.

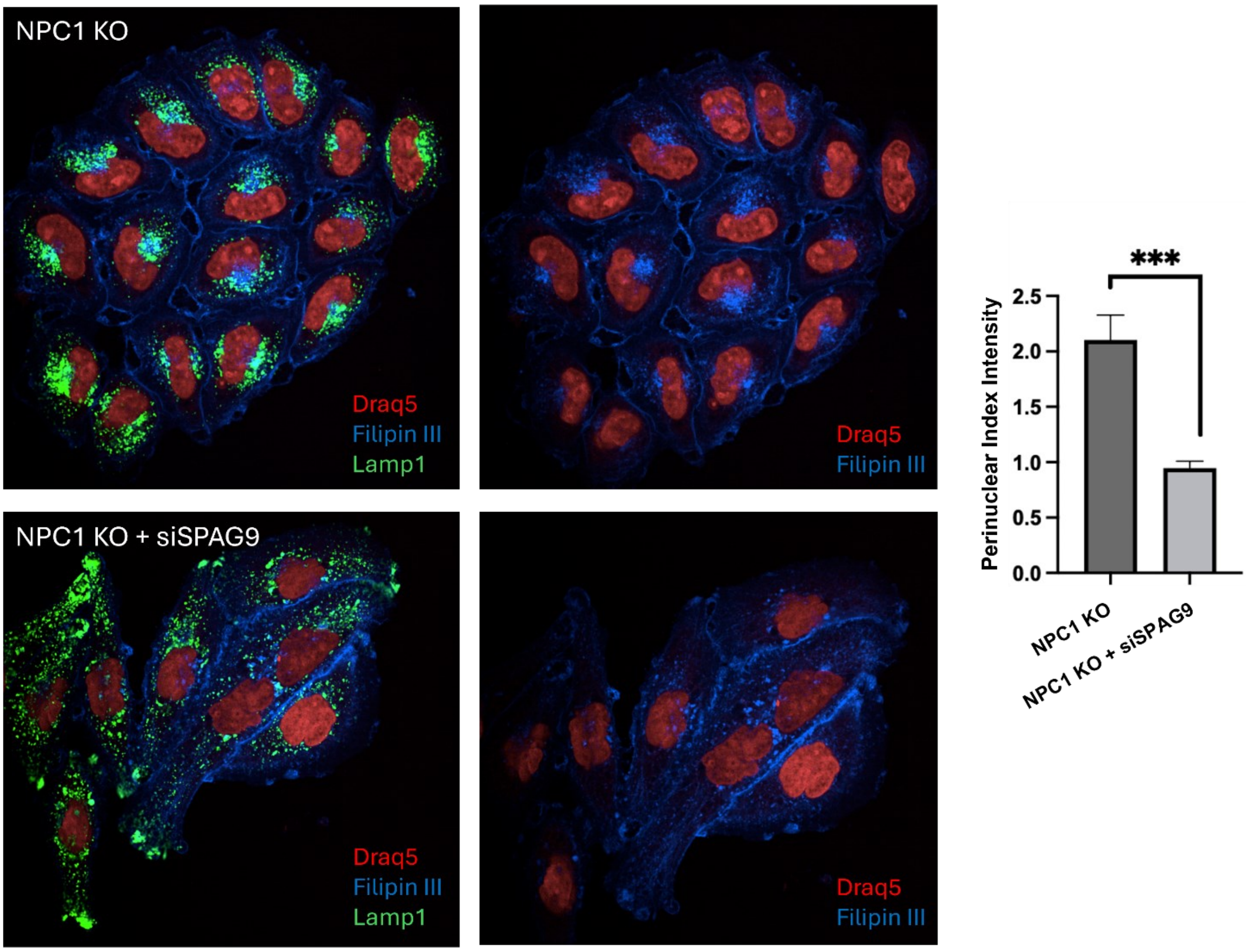


**Fig. S7. The depletion of SPAG9 using siRNAs drastically reduces lysosomal storage around the nucleus in (bottom row) mutant HeLa cells (namely, NPC1 KO + siSPAG9) with respect to the (top row) control HeLa NPC1 KO cells in high-content confocal images.** Cells are stained using Lamp1 (green) to detect lysosomes, Filipin III (blue) to detect cholesterol, and Draq5 (red) to detect nuclei. For the quantification of the lysosomal distribution inside the cell, the cytoplasm is segmented into two areas (perinuclear and peripheral). The perinuclear index intensity on the right reports the ratio between the perinuclear and peripheral mean intensities related to Lamp1. Higher values mean greater perinuclear accumulation of lysosomes. A redistribution of lysosomes near the plasma membrane occurs in the cells after treatment.

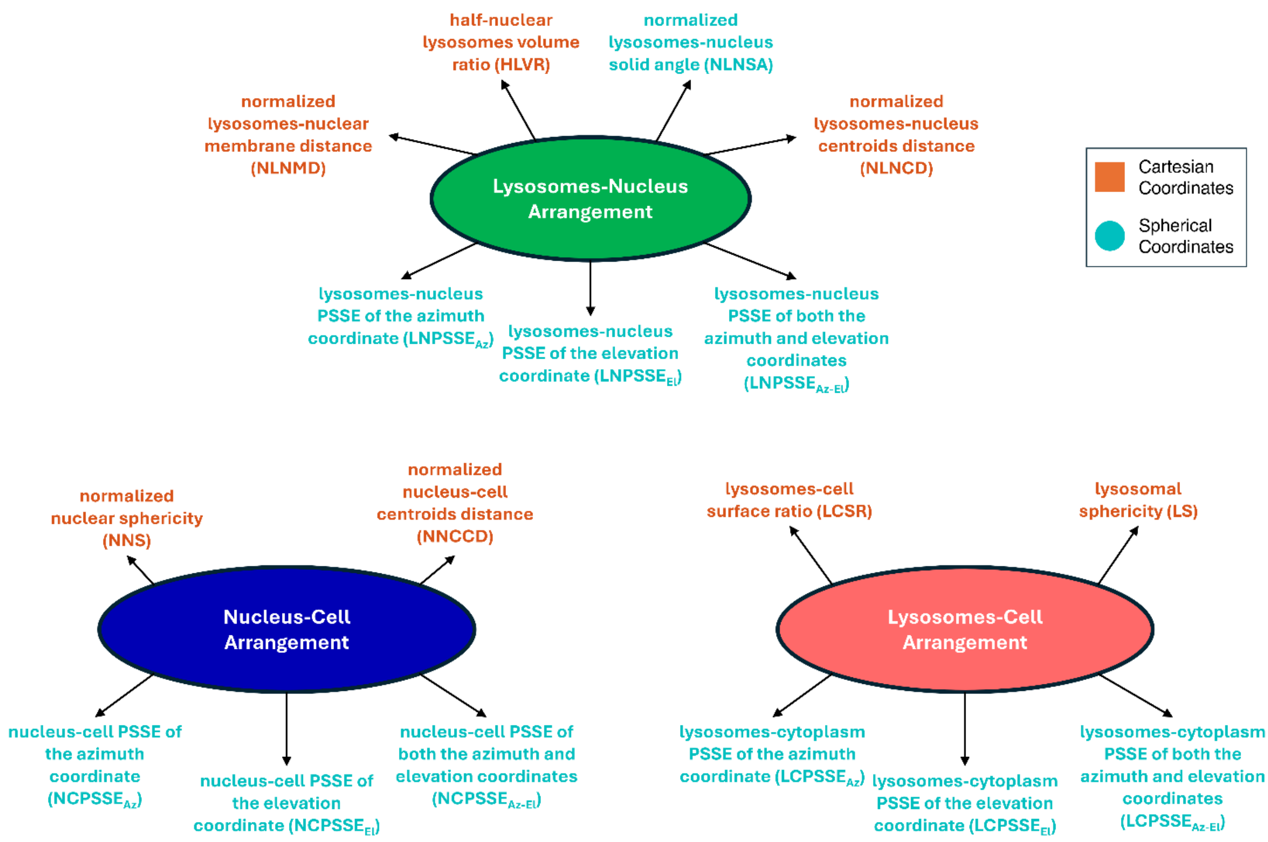


**Fig. S8. Categorization into biomarkers of the 3D morphometric features used to quantitatively characterize the 3D RI tomograms of NPC healthy and diseased cells.** Features can be computed in a cartesian or a spherical coordinate reference system, and they can be categorized into three classes of biomarkers according to the specific intracellular arrangement they describe.

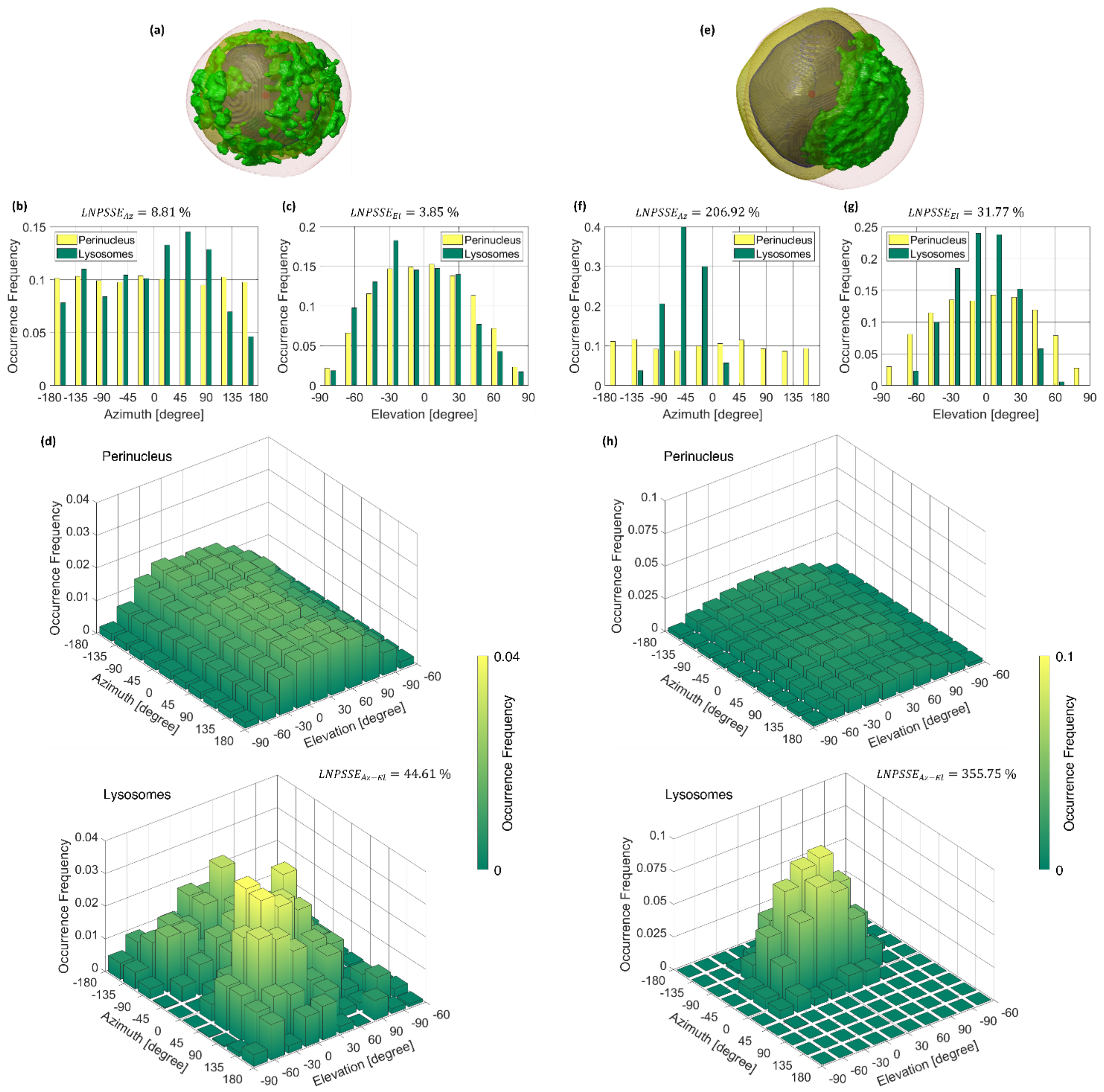


**Fig. S9. Lysosomes-nucleus feature set based on a spherical coordinate system computed from the 3D RI tomogram of a suspended (a-d) HeLa WT cell and (e-g) HeLa NPC1 KO cell. (a,e)** Isolevels representation of the cell (red) and its nucleus (blue), perinucleus (yellow), and LVC (green). The coordinate reference system is centered in the nucleus centroid (red dot). **(b,f)** Normalized histogram of the perinucleus azimuth coordinates (yellow) and the LVC azimuth coordinates (green). **(c,g)** Normalized histogram of the perinucleus elevation coordinates (yellow) and the LVC elevation coordinates (green). **(d,h)** Normalized bivariate histogram of both the azimuth and elevation coordinates about the perinucleus (top) and the LVC (bottom). In (b-d) and (f-h), the LNPSSE values are reported.

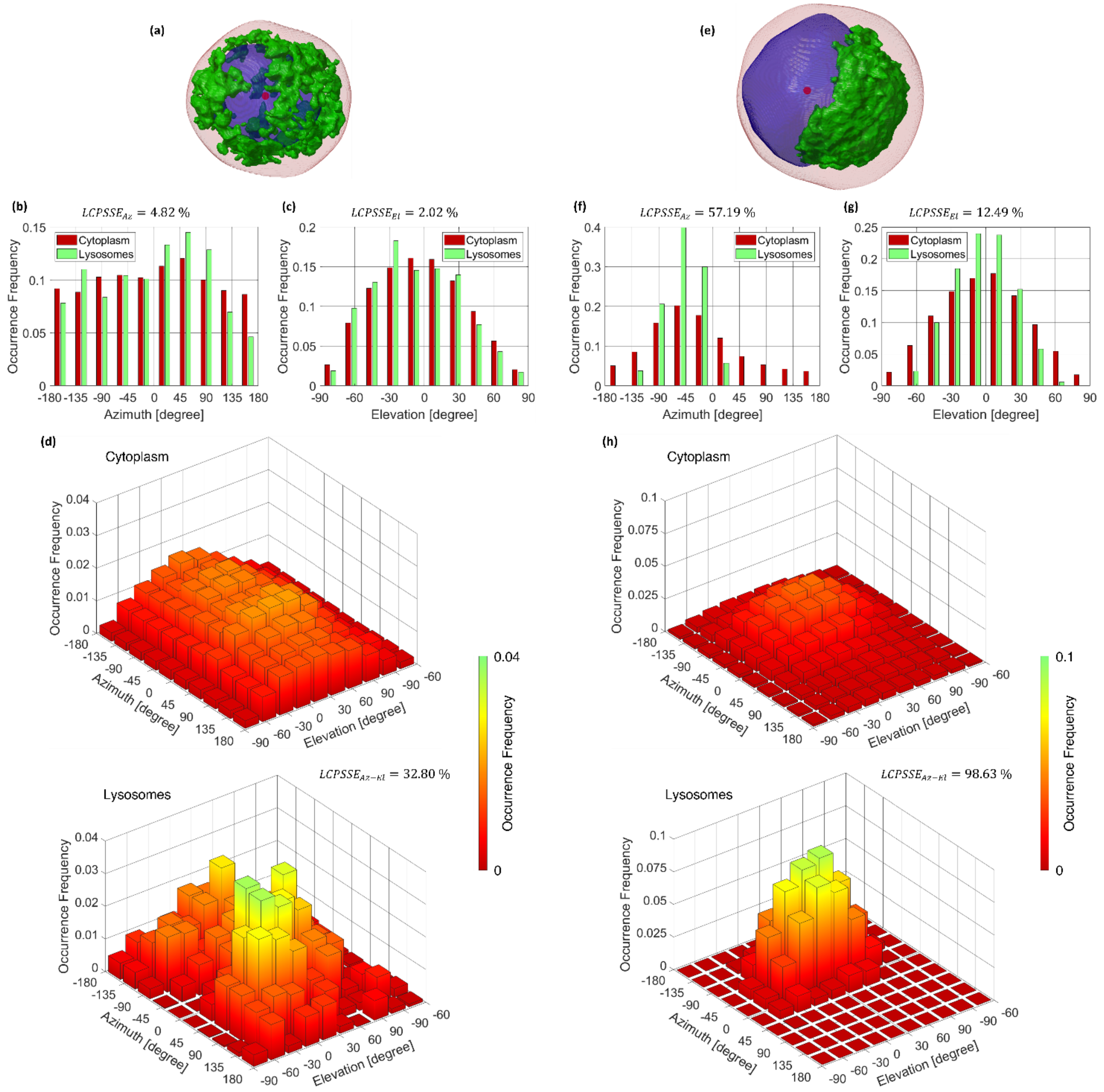


**Fig. S10. Lysosomes-cytoplasm feature set based on a spherical coordinate system computed from the 3D RI tomogram of a suspended (a-d) HeLa WT cell and (e-g) HeLa NPC1 KO cell. (a,e)** Isolevels representation of the nucleus (blue), the LVC (green), and the cytoplasm (red), i.e. the whole cell without the nucleus. The coordinate reference system is centered in the nucleus centroid (red dot). **(b,f)** Normalized histogram of the cytoplasm azimuth coordinates (red) and the LVC azimuth coordinates (green). **(c,g)** Normalized histogram of the cytoplasm elevation coordinates (red) and the LVC elevation coordinates (green). **(d,h)** Normalized bivariate histogram of both the azimuth and elevation coordinates about the cytoplasm (top) and the LVC (bottom). In (b-d) and (f-h), the LCPSSE values are reported.

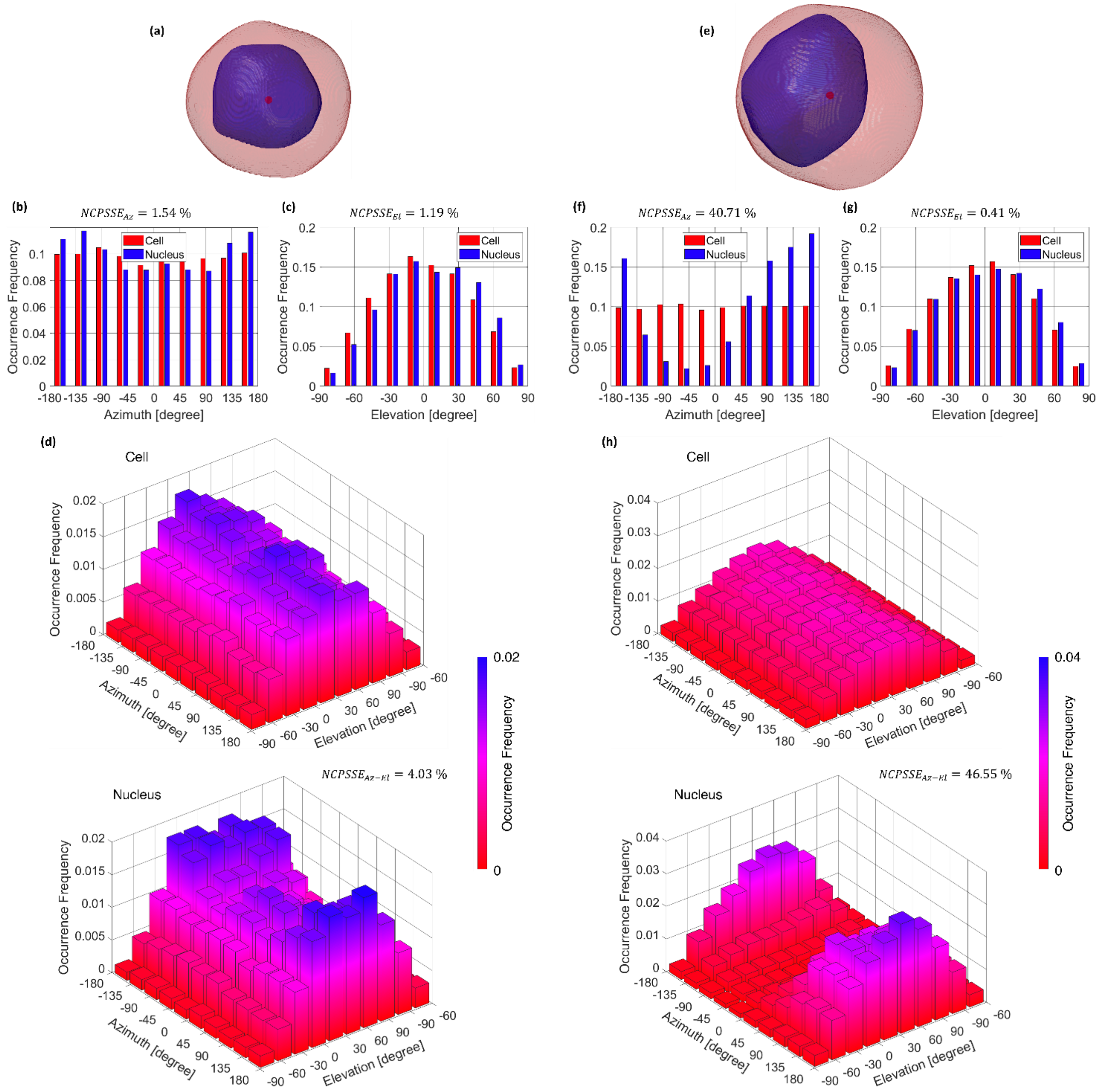


**Fig. S11. Nucleus-cell feature set based on a spherical coordinate system computed from the 3D RI tomogram of a suspended (a-d) HeLa WT cell and (e-g) HeLa NPC1 KO cell. (a,e)** Isolevels representation of the cell (red) and its nucleus (blue). The coordinate reference system is centered in the cell centroid (red dot). **(b,f)** Normalized histogram of the cell azimuth coordinates (red) and the nucleus azimuth coordinates (blue). **(c,g)** Normalized histogram of the cell elevation coordinates (red) and the nucleus elevation coordinates (blue). **(d,h)** Normalized bivariate histogram of both the azimuth and elevation coordinates about the cell (top) and the nucleus (bottom). In (b-d) and (f-h), the NCPSSE values are reported.

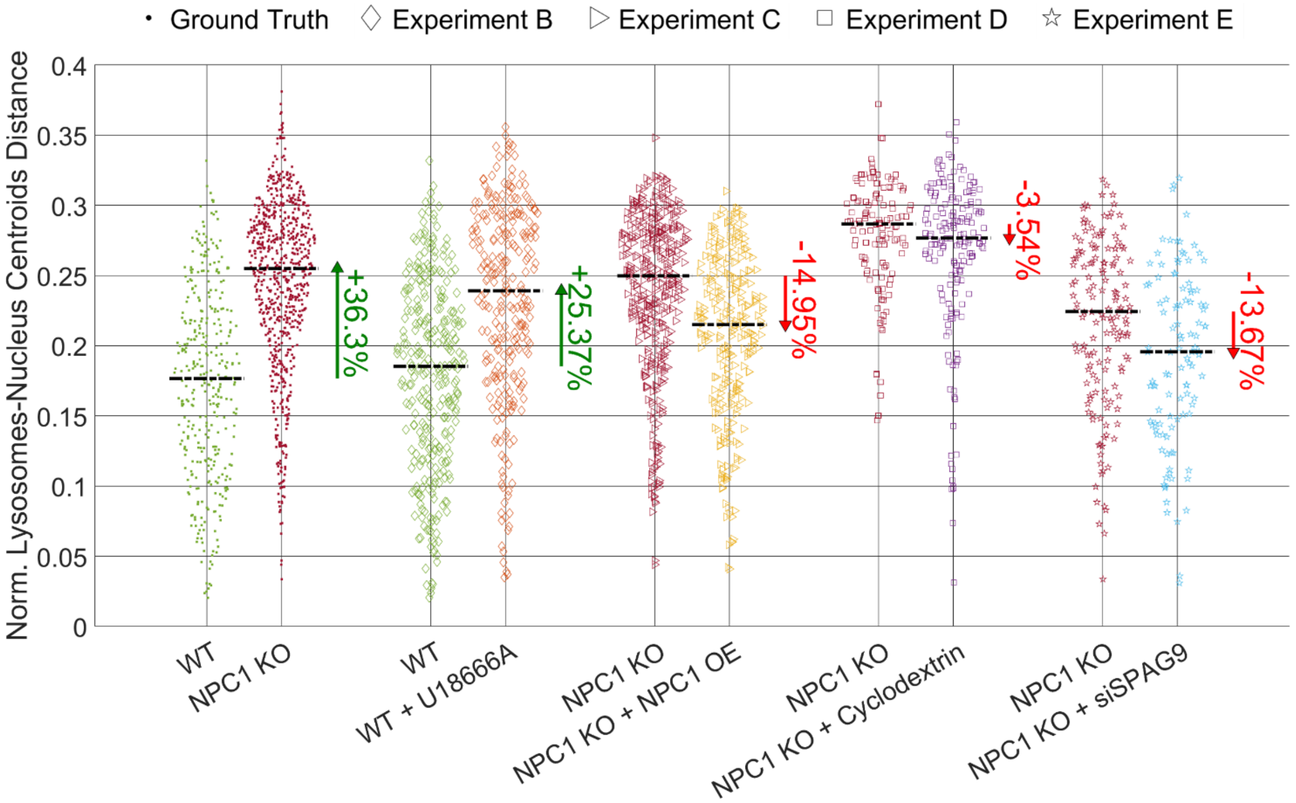


**Fig. S12. Comparison between the NLNCD swarm charts about the ground truth experiment (WT vs. NPC1 KO) and other four experiments (B, C, D, and E) based on specific treatments applied to the HeLa cells to quantitatively evaluate the NPC disease by HTFC.** Black dashed lines are the median values. For each experiment, the PV values are reported (green if positive, red if negative).

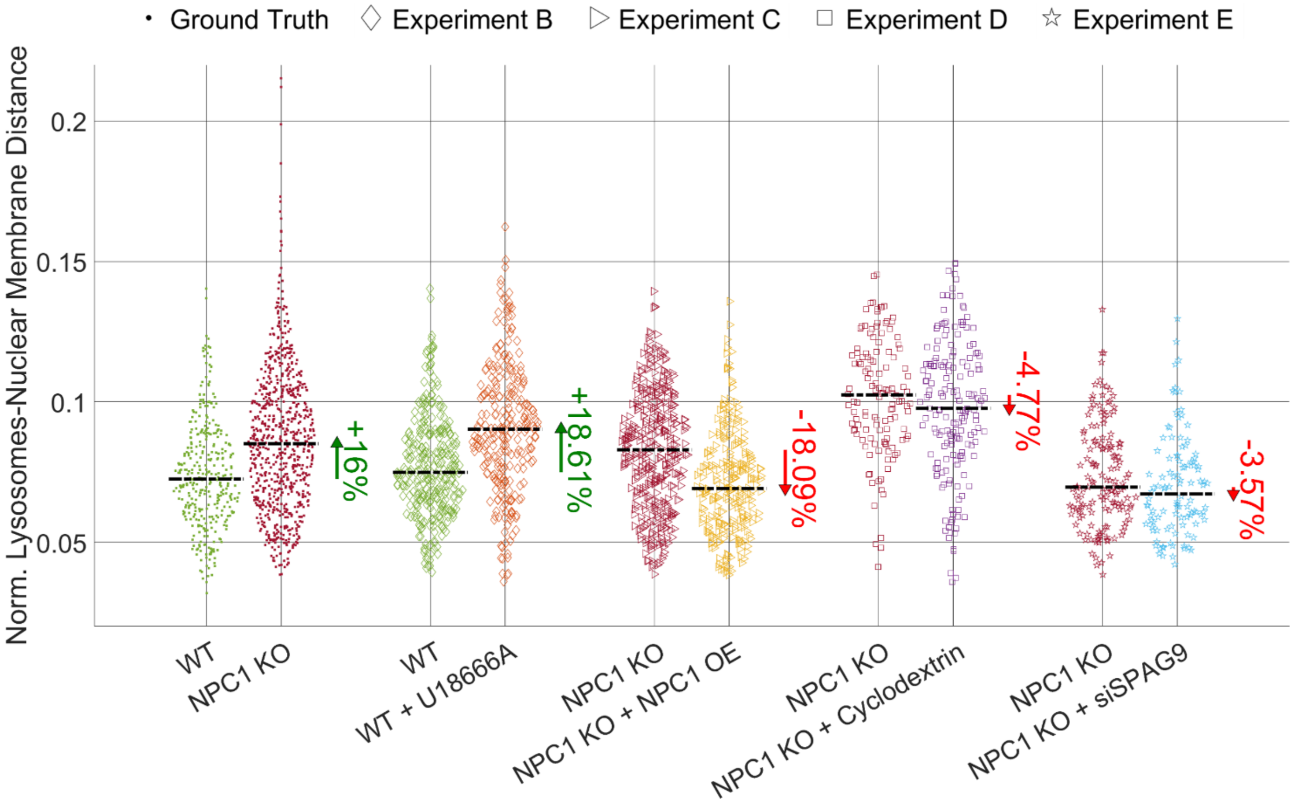


**Fig. S13. Comparison between the NLNMD swarm charts about the ground truth experiment (WT vs. NPC1 KO) and other four experiments (B, C, D, and E) based on specific treatments applied to the HeLa cells to quantitatively evaluate the NPC disease by HTFC.** Black dashed lines are the median values. For each experiment, the PV values are reported (green if positive, red if negative).

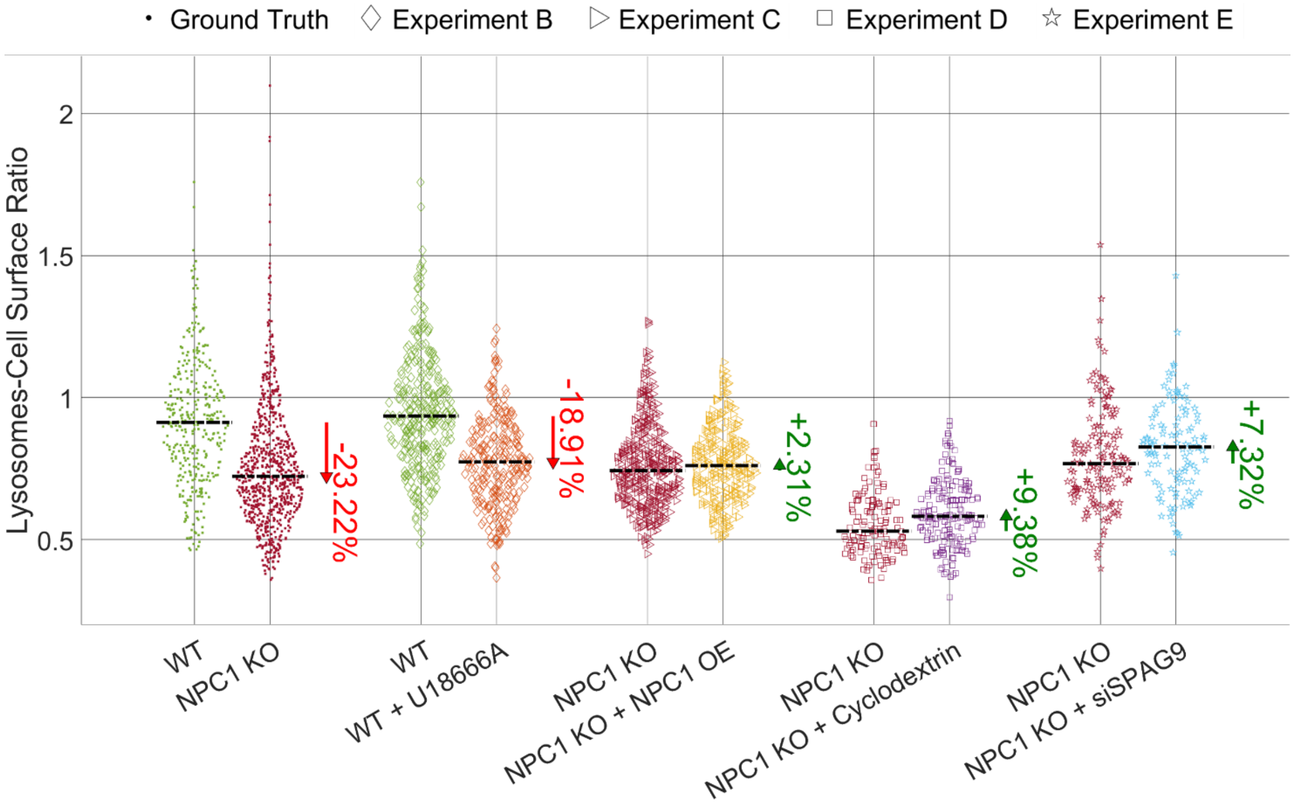


**Fig. S14. Comparison between the LCSR swarm charts about the ground truth experiment (WT vs. NPC1 KO) and other four experiments (B, C, D, and E) based on specific treatments applied to the HeLa cells to quantitatively evaluate the NPC disease by HTFC.** Black dashed lines are the median values. For each experiment, the PV values are reported (green if positive, red if negative).

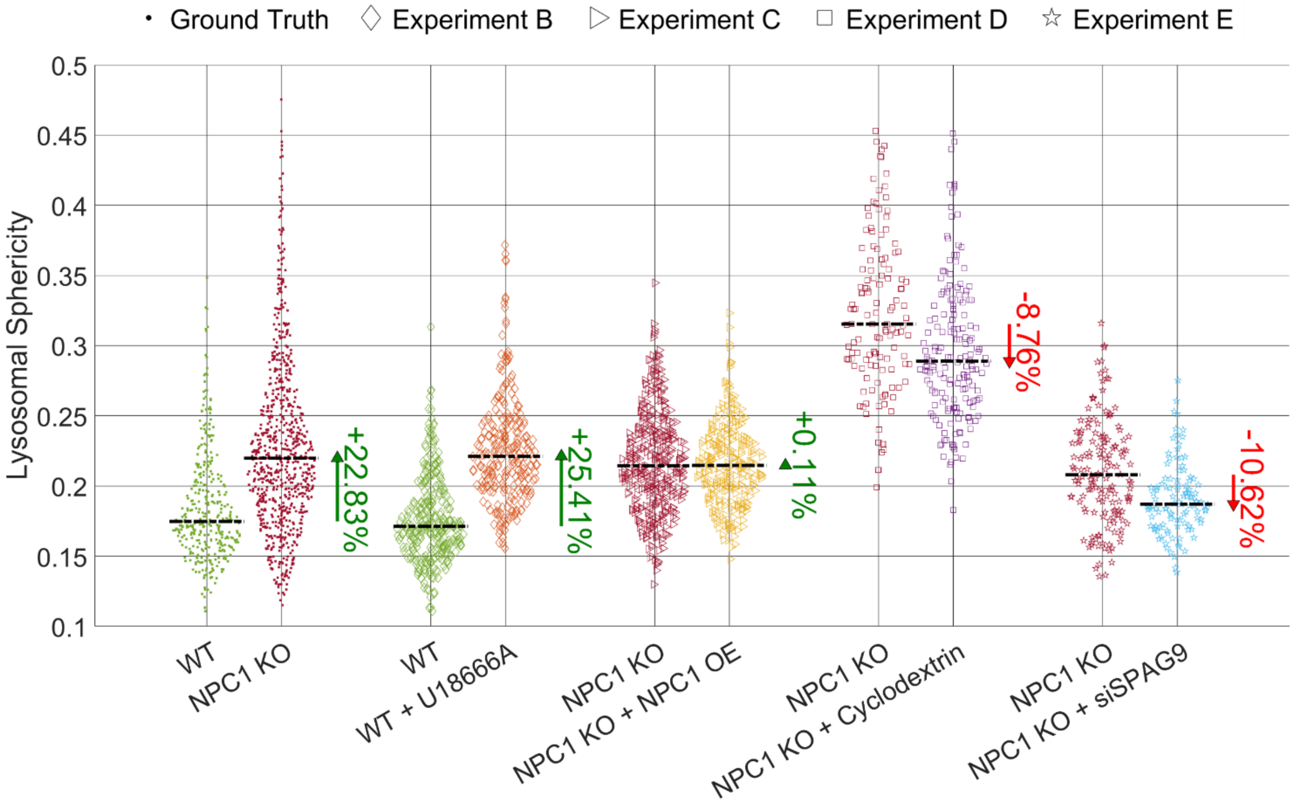


**Fig. S15. Comparison between the LS swarm charts about the ground truth experiment (WT vs. NPC1 KO) and other four experiments (B, C, D, and E) based on specific treatments applied to the HeLa cells to quantitatively evaluate the NPC disease by HTFC.** Black dashed lines are the median values. For each experiment, the PV values are reported (green if positive, red if negative).

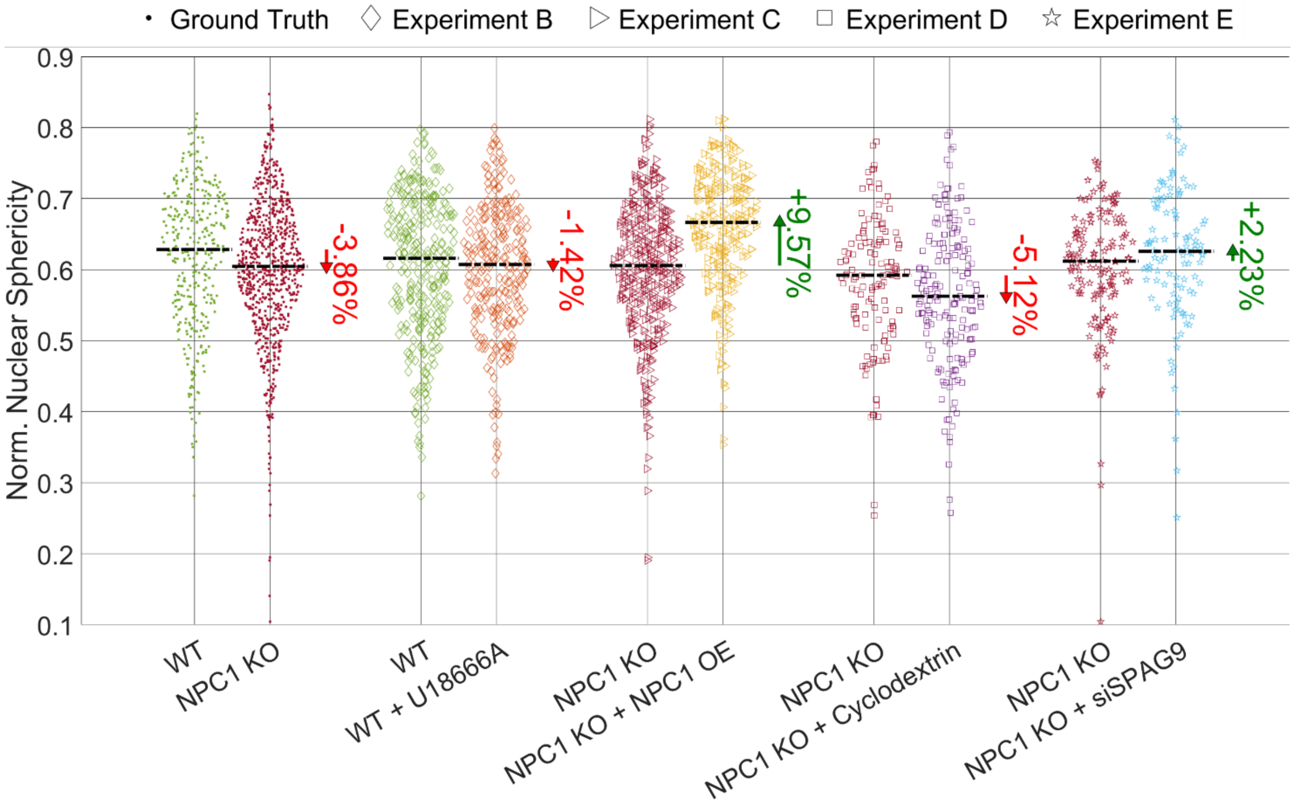


**Fig. S16. Comparison between the NNS swarm charts about the ground truth experiment (WT vs. NPC1 KO) and other four experiments (B, C, D, and E) based on specific treatments applied to the HeLa cells to quantitatively evaluate the NPC disease by HTFC.** Black dashed lines are the median values. For each experiment, the PV values are reported (green if positive, red if negative).

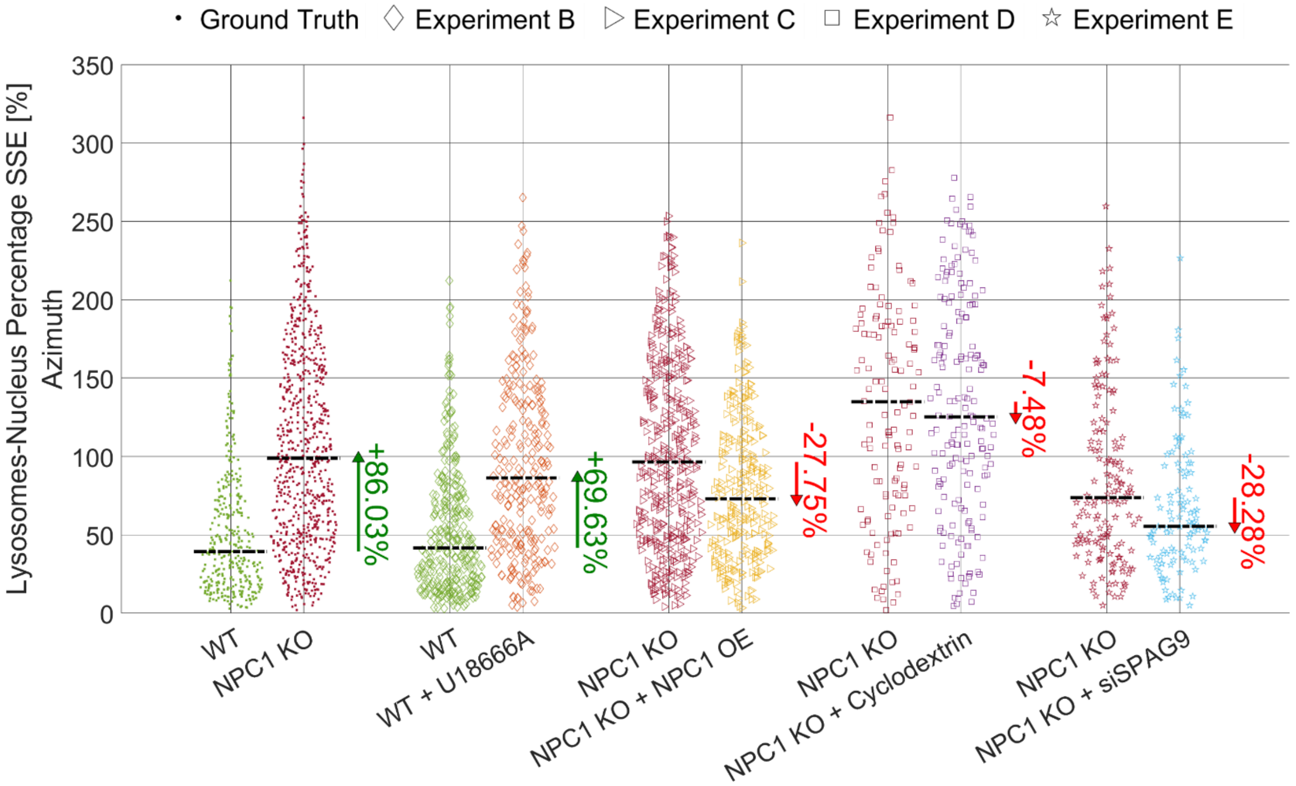


**Fig. S17. Comparison between the LNPSSE$_{Az}$ swarm charts about the ground truth experiment (WT vs. NPC1 KO) and other four experiments (B, C, D, and E) based on specific treatments applied to the HeLa cells to quantitatively evaluate the NPC disease by HTFC.** Black dashed lines are the median values. For each experiment, the PV values are reported (green if positive, red if negative).

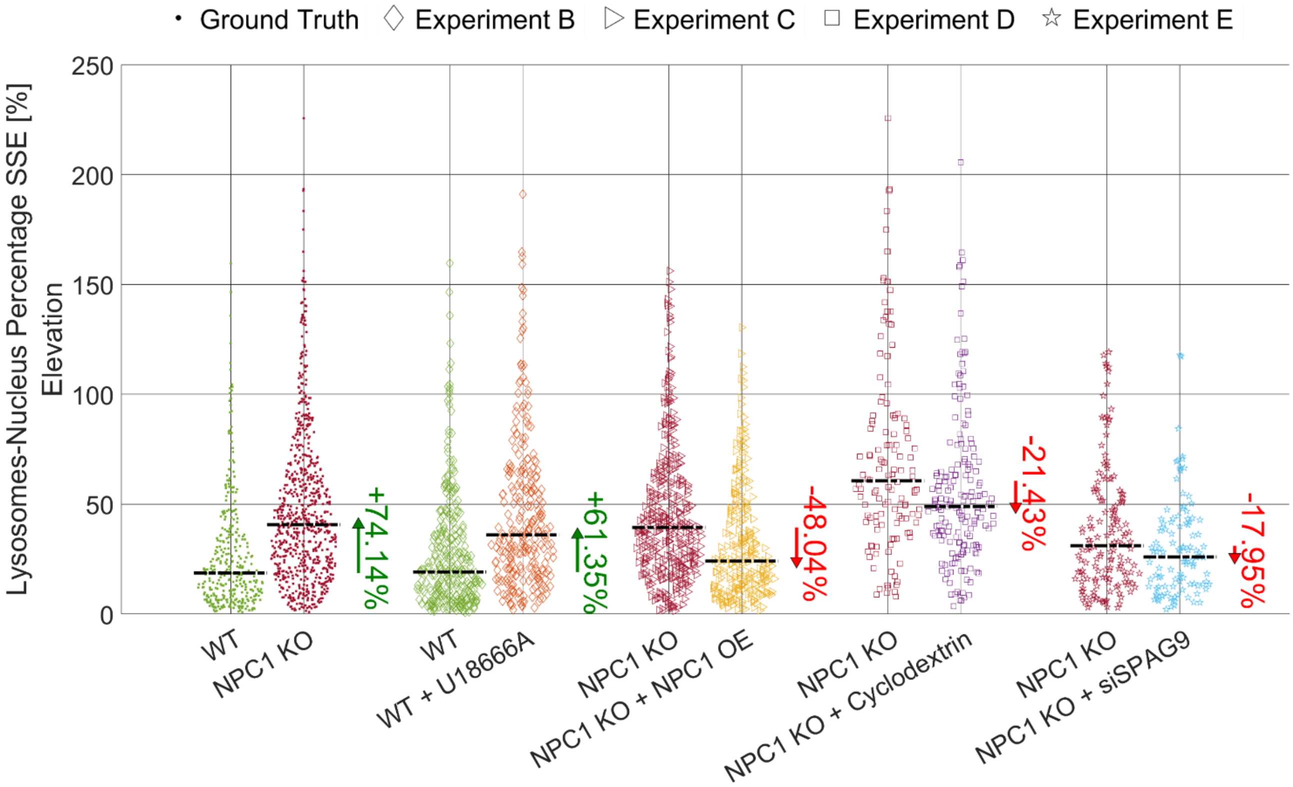


**Fig. S18. Comparison between the $LNPSSE_{El}$ swarm charts about the ground truth experiment (WT vs. NPC1 KO) and other four experiments (B, C, D, and E) based on specific treatments applied to the HeLa cells to quantitatively evaluate the NPC disease by HTFC.** Black dashed lines are the median values. For each experiment, the PV values are reported (green if positive, red if negative).

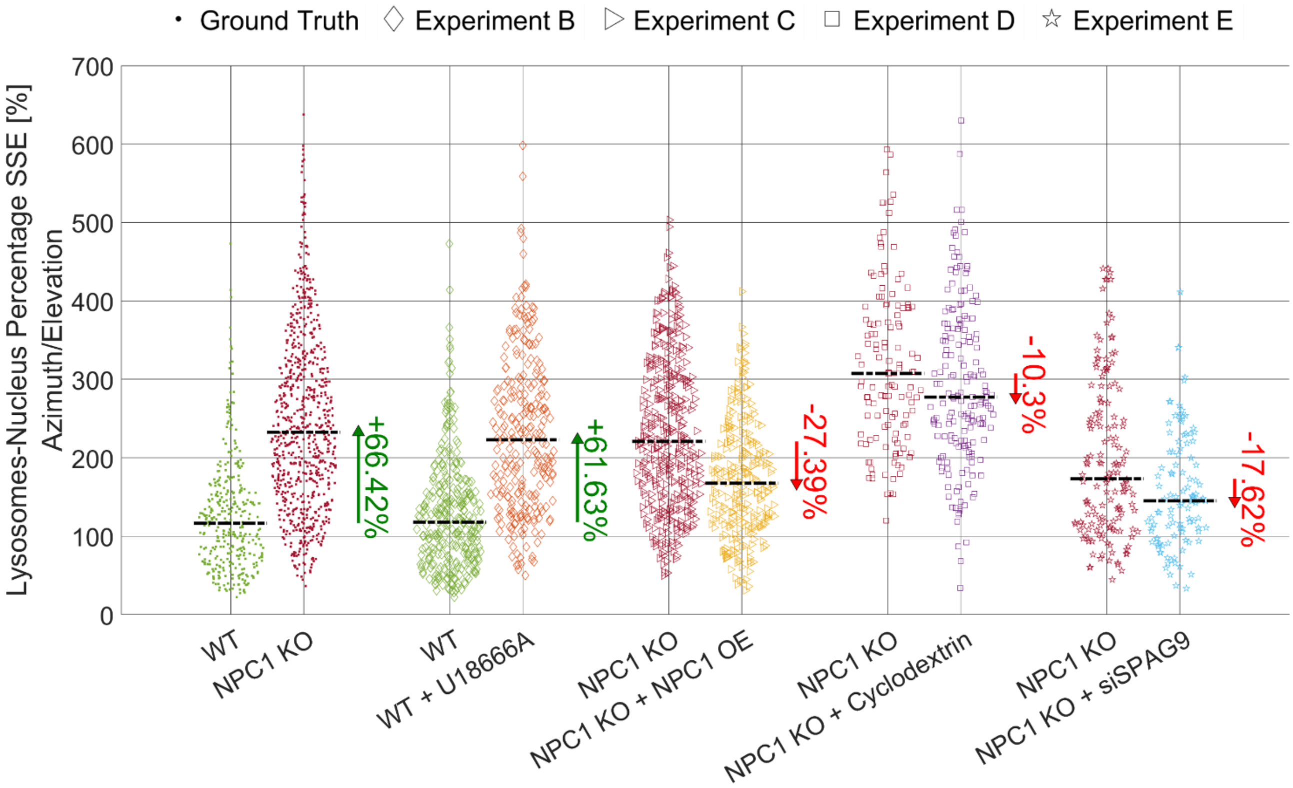


**Fig. S19. Comparison between the $LNPSSE_{Az-El}$ swarm charts about the ground truth experiment (WT vs. NPC1 KO) and other four experiments (B, C, D, and E) based on specific treatments applied to the HeLa cells to quantitatively evaluate the NPC disease by HTFC.** Black dashed lines are the median values. For each experiment, the PV values are reported (green if positive, red if negative).

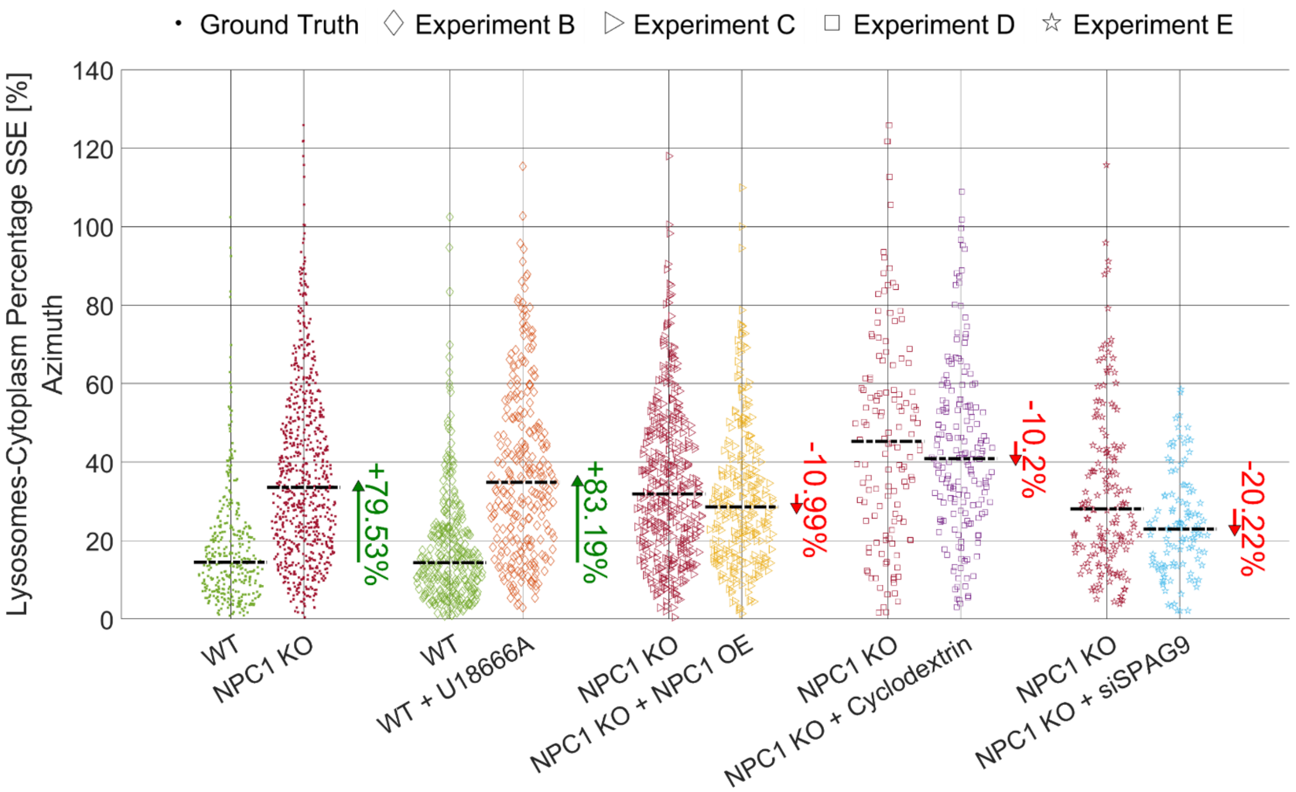


**Fig. S20. Comparison between the $LCPSSE_{Az}$ swarm charts about the ground truth experiment (WT vs. NPC1 KO) and other four experiments (B, C, D, and E) based on specific treatments applied to the HeLa cells to quantitatively evaluate the NPC disease by HTFC.** Black dashed lines are the median values. For each experiment, the PV values are reported (green if positive, red if negative).

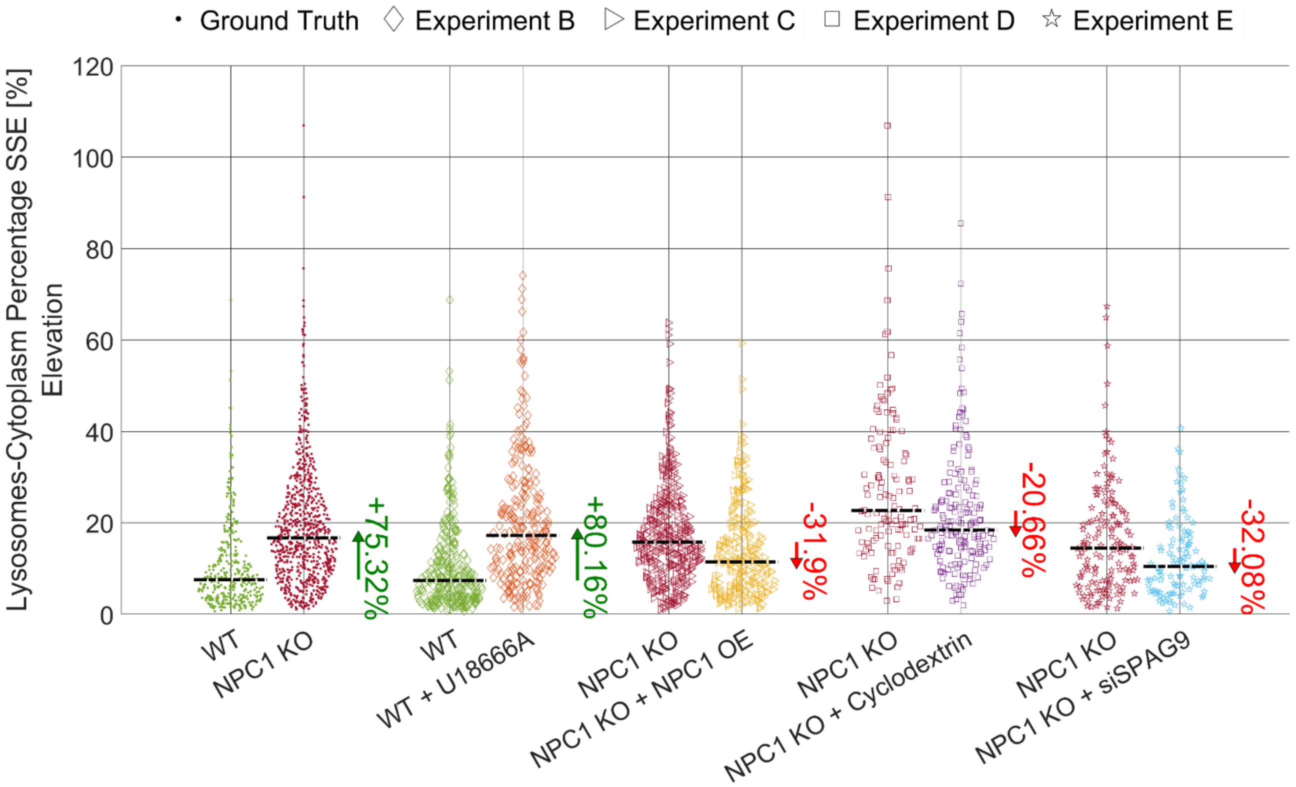


**Fig. S21. Comparison between the $LCPSSE_{El}$ swarm charts about the ground truth experiment (WT vs. NPC1 KO) and other four experiments (B, C, D, and E) based on specific treatments applied to the HeLa cells to quantitatively evaluate the NPC disease by HTFC.** Black dashed lines are the median values. For each experiment, the PV values are reported (green if positive, red if negative).

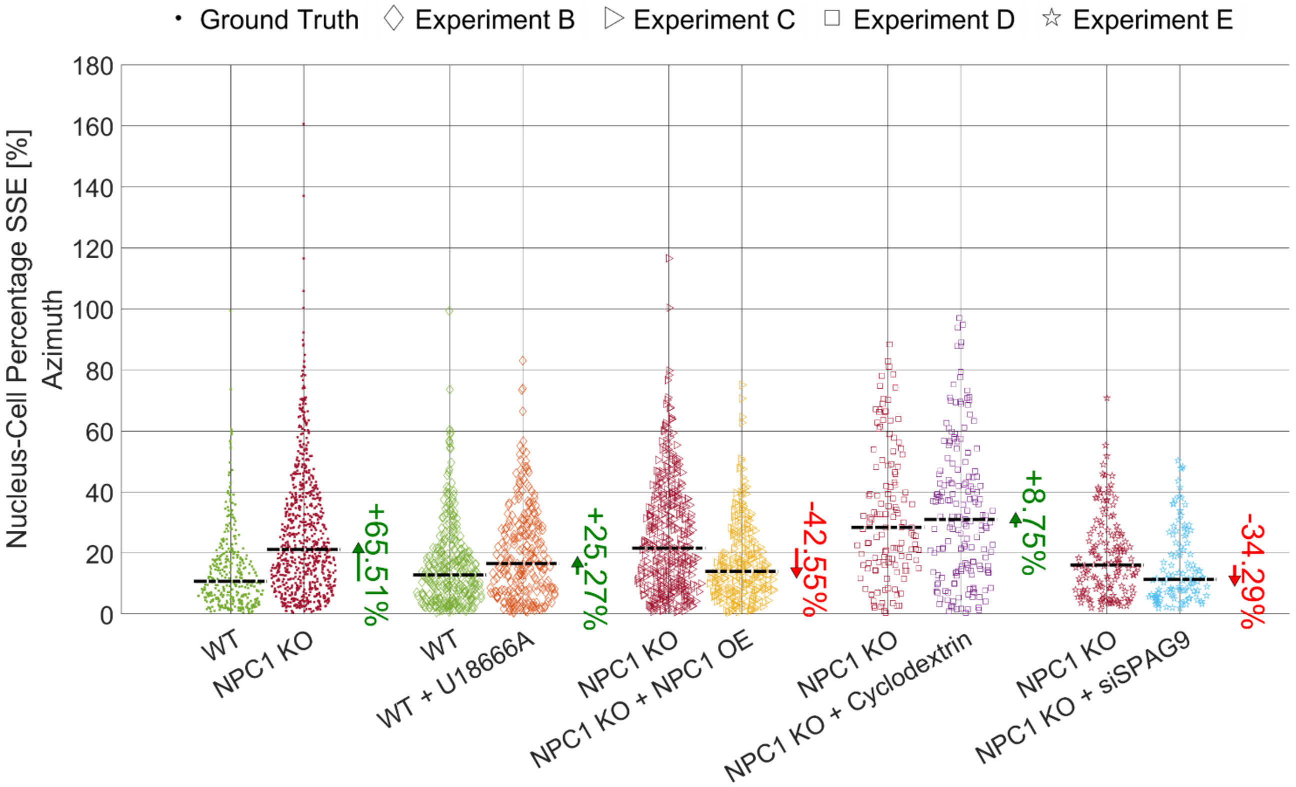


**Fig. S22. Comparison between the $NCPSSE_{Az}$ swarm charts about the ground truth experiment (WT vs. NPC1 KO) and other four experiments (B, C, D, and E) based on specific treatments applied to the HeLa cells to quantitatively evaluate the NPC disease by HTFC.** Black dashed lines are the median values. For each experiment, the PV values are reported (green if positive, red if negative).

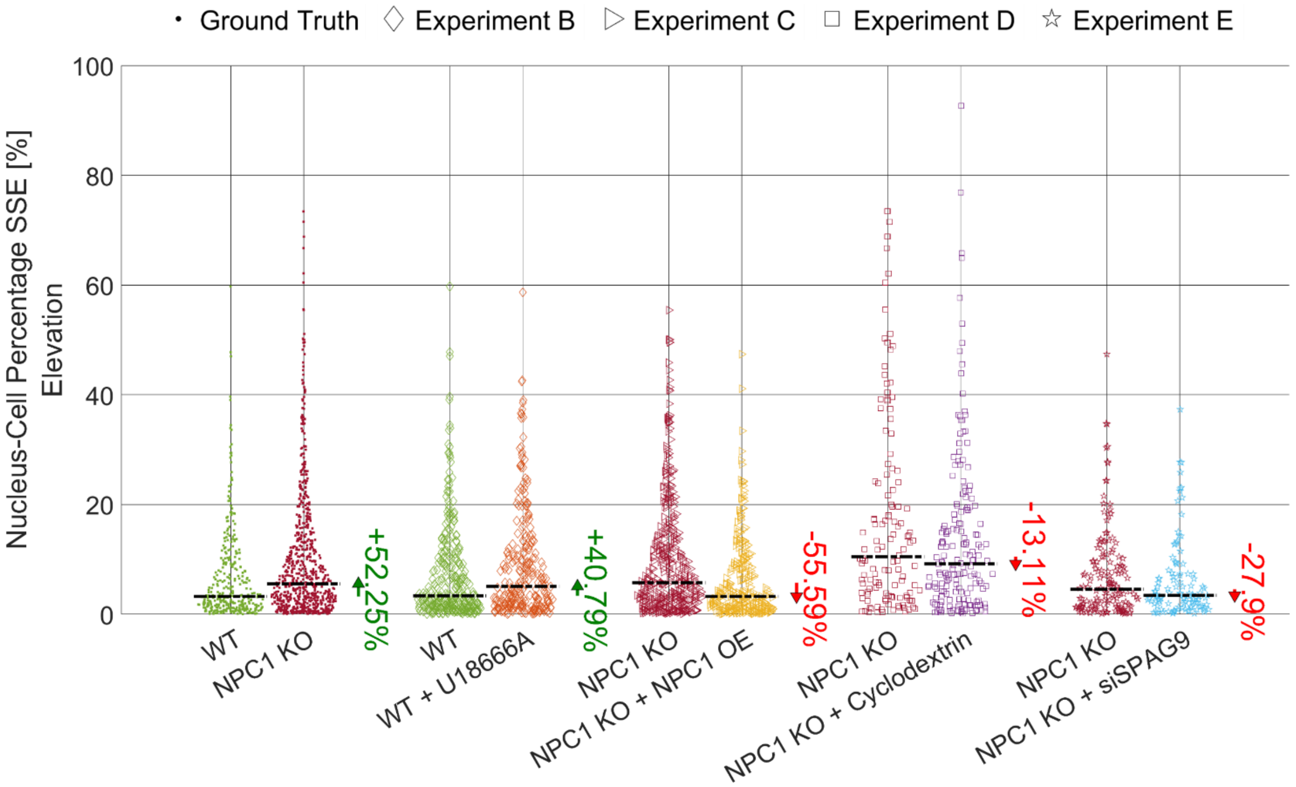


**Fig. S23. Comparison between the $NCPSSE_{El}$ swarm charts about the ground truth experiment (WT vs. NPC1 KO) and other four experiments (B, C, D, and E) based on specific treatments applied to the HeLa cells to quantitatively evaluate the NPC disease by HTFC.** Black dashed lines are the median values. For each experiment, the PV values are reported (green if positive, red if negative).

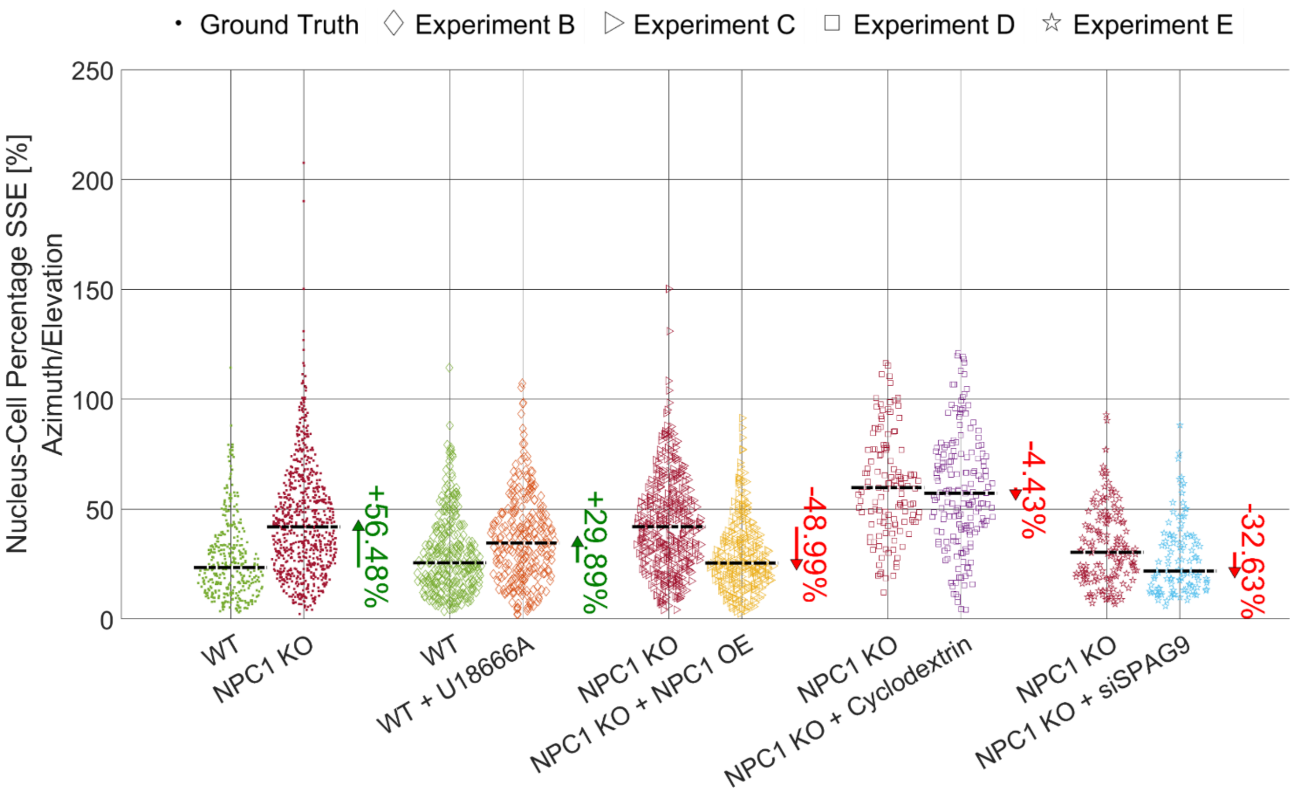


**Fig. S24. Comparison between the NCPSSE$_{Az\text{-}El}$ swarm charts about the ground truth experiment (WT vs. NPC1 KO) and other four experiments (B, C, D, and E) based on specific treatments applied to the HeLa cells to quantitatively evaluate the NPC disease by HTFC.** Black dashed lines are the median values. For each experiment, the PV values are reported (green if positive, red if negative).

**Table S1.** Overall dataset of HeLa cells collected in 5 different experiments by HTFC.

| *Experiment* / *Cell Line* | A | B | C | D | E | Total |
|---|---|---|---|---|---|---|
| **WT** | 53 | 305 | - | - | - | 358 |
| **NPC1 KO** | 107 | - | 381 | 128 | 150 | 766 |
| **WT + U18666A** | - | 262 | - | - | - | 262 |
| **NPC1 KO + NPC1 OE** | - | - | 268 | - | - | 268 |
| **NPC1 KO + Cyclodextrin** | - | - | - | 177 | - | 177 |
| **NPC1 KO + siSPAG9** | - | - | - | - | 108 | 108 |
| | | | | | | 1939 |

**Table S2.** List of morphometric biomarkers to quantitatively characterize the 3D RI tomograms of NPC healthy and diseased cells, along with corresponding acronym and value range.

| Feature | Acronym | Range |
|---|---|---|
| half-nuclear lysosomes volume ratio | HLVR | [0.5,1] |
| normalized lysosomes-nucleus solid angle | NLNSA | [0,1] |
| normalized lysosomes-nucleus centroids distance | NLNCD | [0,2] |
| normalized lysosomes-nuclear membrane distance | NLNMD | [0,2] |
| lysosomes-cell surface ratio | LCSR | [0,∞] |
| lysosomal sphericity | LS | [0,1] |
| normalized nuclear sphericity | NNS | [0,1] |
| normalized nucleus-cell centroids distance | NNCCD | [0,2] |
| lysosomes-nucleus PSSE of the azimuth coordinate | $\mathrm{LNPSSE}_{\mathrm{Az}}$ | [0,100] % |
| lysosomes-nucleus PSSE of the elevation coordinate | $\mathrm{LNPSSE}_{\mathrm{El}}$ | [0,100] % |
| lysosomes-nucleus PSSE of both the azimuth and elevation coordinates | $\mathrm{LNPSSE}_{\mathrm{Az-El}}$ | [0,100] % |
| lysosomes-cytoplasm PSSE of the azimuth coordinate | $\mathrm{LCPSSE}_{\mathrm{Az}}$ | [0,100] % |
| lysosomes-cytoplasm PSSE of the elevation coordinate | $\mathrm{LCPSSE}_{\mathrm{El}}$ | [0,100] % |
| lysosomes-cytoplasm PSSE of both the azimuth and elevation coordinates | $\mathrm{LCPSSE}_{\mathrm{Az-El}}$ | [0,100] % |
| nucleus-cell PSSE of the azimuth coordinate | $\mathrm{NCPSSE}_{\mathrm{Az}}$ | [0,100] % |
| nucleus-cell PSSE of the elevation coordinate | $\mathrm{NCPSSE}_{\mathrm{El}}$ | [0,100] % |
| nucleus-cell PSSE of both the azimuth and elevation coordinates | $\mathrm{NCPSSE}_{\mathrm{Az-El}}$ | [0,100] % |
